# When killers become thieves: trogocytosed PD-1 inhibits NK cells in cancer

**DOI:** 10.1101/2020.06.26.174342

**Authors:** Mohammed S. Hasim, Marie Marotel, Jonathan J. Hodgins, Elisabetta Vulpis, Han-Yun Shih, Amit Scheer, Olivia MacMillan, Fernando G. Alonso, Kelly P. Burke, David P. Cook, Maria Teresa Petrucci, Angela Santoni, Padraic G. Fallon, Arlene H. Sharpe, Giuseppe Sciumè, Andre Veillette, Alessandra Zingoni, Arleigh McCurdy, Michele Ardolino

## Abstract

Leucocytes often perform trogocytosis, the process by which cells acquire parts of the plasma membrane from interacting cells. Accumulating evidence indicates that trogocytosis modulates immune responses, but the underlying molecular mechanisms are unclear. Here, using two mouse models of leukemia, we found that cytotoxic lymphocytes perform trogocytosis at high rates with tumor cells. While performing trogocytosis, both Natural Killer and CD8^+^ T cells acquire the checkpoint receptor PD-1 from leukemia cells. In vitro and in vivo investigation revealed that PD-1 protein found on the surface of Natural Killer cells, rather than being endogenously expressed, was derived entirely from leukemia cells. Mechanistically, SLAM receptors were essential for PD-1 trogocytosis. PD-1 acquired via trogocytosis actively suppressed anti-tumor immunity, as revealed by the positive outcome of PD-1 blockade in PD-1-deficient mice. PD-1 trogocytosis was corroborated in patients with clonal plasma cell disorders, where Natural Killer cells that stained for PD-1 also stained for tumor cell markers. Our results, in addition to shedding light on a previously unappreciated mechanism underlying the presence of PD-1 on Natural Killer and cytotoxic T cells, reveal the immune-regulatory effect of membrane transfer occurring when immune cells contact tumor cells.

**Once sentence summary:** Natural Killer cells are inhibited by PD-1 acquired from the surface of tumor cells via trogocytosis.

## Introduction

During trogocytosis immune cells acquire parts of the membrane of cells they interact with (*1, 2*). First characterized in ⍺β-T cells (*3–8*), it later became clear that virtually all immune cells perform trogocytosis (*7, 9–16*). This intercellular transfer of membranes results in the acquisition of proteins that would otherwise not be endogenously expressed by the cell performing trogocytosis, as in the case of NK cells that acquire viral proteins from infected cells (*17, 18*), or cancer antigens from tumor cells (*19*). Proteins transferred via trogocytosis are functional and influence the response of the accepting cell (*11, 16, 18, 20-24*). The pathophysiological relevance of trogocytosis is underscored by the high extent that immune cells perform it in the context of infections (*25, 26*), autoimmune diseases (*27*), and cancer (*23, 28, 29*). Natural Killer (NK) cells are important mediators of the response against intracellular pathogens and tumors (*30–32*) and have been amongst the first immune cells shown to perform trogocytosis (*10–12*). Trogocytosis has been reported to contribute to the negative regulation of NK cell responses in different contexts. For example, acquisition of m157 or NKG2D ligands results in sustained and unproductive crosslinking of activating receptors leading to NK cell anergy (*18, 33, 34*), but also promotes NK fratricide (*34, 35*). On the other hand, acquisition of MHC molecules from target cells engaged Ly49 receptors in *cis*, sustaining inhibitory signaling that dampened NK cell activation (*11*). Finally, trogocytosis of HLA-G from cancer cells resulted in the generation of NK cells with suppressive properties (*36*).

We recently reported that NK cells are suppressed by the checkpoint receptor PD-1 and contribute to the therapeutic efficacy of PD-1/L1 blockade in mouse models of cancer (*37*). These results, corroborated by others (*38–43*), were at least partially confuted by findings that murine and human NK cells fail to endogenously express *Pdcd1* mRNA or PD-1 protein (*44*). In light of our results indicating that PD-1 is found on the surface of NK cells, and considering the high trogocytosis activity of NK cells, we propose that NK cells acquire PD-1 directly from tumor cells. Mechanistic experiments corroborated our hypothesis and revealed that SLAM receptors were essential for PD-1 trogocytosis. Functionally, trogocytosed PD-1 suppressed NK cell mediated cancer immunosurveillance. Finally, analysis of NK cells in patients with clonal plasma cell disorders suggests that PD-1 trogocytosis occurs in cancer patients. Altogether, our data shed light on a new mechanism that regulates NK cell function via acquisition of PD-1 from tumor cells.

### Materials and Methods. Mice and in vivo procedures

Mice were maintained at the University of Ottawa. *Pdcd1* knockout mice (B6.Cg-Pdcd1tm1.1Shr/J)(*45*) were purchased from The Jackson Laboratory and crossed with C57BL/6J mice purchased from The Jackson Laboratory to obtain *Pdcd1* heterozygous mice. Heterozygous mice were bred to obtain *Pdcd1*^+/+^ and *Pdcd1^-/-^* littermates. *Ncr1^+/Cre^* mice (*46*) were kindly gifted by Dr. Vivier (INSERM, Marseille, France) and crossed with *Cd274^fl/fl^* mice (*47*), kindly gifted by Dr. Fallon (Trinity College, Dublin, Ireland). Mice were then crossed with *Pdcd1^-/-^* mice. *Cd274^-/-^* mice (*48*) were obtained from Dr. Sharpe (Harvard Medical School, Boston, MA). SLAM-ko mice (*49*) were donated by Dr. Veillette (Institut de recherches cliniques de Montréal, Montréal, QC). *Itgal1^-/-^* mice (*50*) were purchased from The Jackson Laboratory. *Klrk1^-/-^* mice (*51*) and B6 Cd45.1 mice were kindly gifted by Dr. Raulet (University of California, Berkeley, Berkeley, CA). NCG mice were purchased from Charles Rivers Laboratories. For all experiments, sex-matched (both males and females) and age-matched (7 to 18 weeks old) mice were used.

For subcutaneous injections, tumor cells were resuspended in 100 µl PBS and injected in the left flank. Tumors were collected when tumor volume was approximately 300 mm^3^. In some experiments, 0.5x10^6^ tumor cells were resuspended in 100 µl Growth Factor Reduced Matrigel (BD) and injected in both the left and right flank of the same mouse.

Tumor outgrowth of parental or PD-1-deficient RMA-S-*Pdl1* cells was assessed in *Pdcd1^-/-^* or NCG mice injected with 0.1 x10^6^ tumor cells resuspended in 100 µl Matrigel.

For immunotherapy experiments, 0.5x10^6^ tumor cells were resuspended in 100 µl Matrigel mixed with 20 µg of anti-PD-1 (RMP1-14) or control antibody (1–1) (both by Leinco). In some experiments, Fc-silent RMP-14 (*52*) was used.

When indicated, mice were depleted of NK cells with i.p. injection of 200 µg of NKR-P1C antibody (PK136, Leinco).

### Cell lines

All cell lines were cultured at 37°C in a humidified atmosphere containing 5% CO_2_ and maintained in RPMI culture medium containing 100 U/ml penicillin, 100 μg/ml streptomycin, 0.2 mg/ml glutamine, 10 μg/ml gentamycin sulfate, 20 mM HEPES, and 5% FCS. Cell line identity was confirmed by flow cytometry when possible, and cells were regularly tested for mycoplasma.

### Ex vivo experiments

Murine splenic NK cells were isolated using the EasySep™ Mouse NK Cell Isolation Kit (StemCell Technologies). In all experiments with isolated NK cells, 1000 U/mL rhIL-2 (NIH BRB Preclinical Repository) was added to the culture medium. In most co-culture experiments NK cells were labelled with Cell Trace Violet proliferation dye (BD Bioscience) and tumor cells with CFSE (Biolegend), or vice-versa. 100,000 NK cells were co-cultured with tumor cells at a 1:1 ratio in 24 well plates in a final volume of 1 mL. When whole spleens were used, 200,000 splenocytes were co-cultured with tumor cells at a 2:1 ratio in 6 well plates, in a final volume of 3 ml.

In ex vivo cytokine stimulation experiments, isolated splenic NK cells were cultured with 10ng/ml or 100ng/ml of IL-15 (Peprotech), 1,000U/ml IL-2, 100ng/ml IL-5 (Peprotech), 100 ng/ml IL-6 (Peprotech), 20ng/ml IL-12 (Peprotech) + 100 ng/ml IL-18 (Leinco), 10ng/ml TGF-β1 (Peprotech), 1,000U/ml Type I IFN (PBL Assay) or 25nM of the glucocorticoid Corticosterone (Sigma) for 3 days.

For transwell experiments (0.4 μm filter, Millipore), co-culture was set up in 6-well plates with a final volume of 3 mL.

In sup transfer experiments, RMA cells were seeded at 200,000 cells/ml and cultured for 3 days. Conditioned media was collected, centrifuged, filtered, diluted 1:1 with fresh media and added to NK cells for 24 hrs.

For membrane dye transfer experiments, NK cells were labelled with CFSE and RMA cells with CellVue Claret FarRed (Sigma-Aldrich). 10,000 NK cells were then co-cultured with RMA cells at a 1:10 ratio in 96-well V-bottom plates with a final volume of 100 µL.

In experiments where ATP production was pharmacologically blocked, NK cells were pre-treated with 50 mM of Sodium Azide (Sigma) for 2hrs or with 13 µM of Antimycin-A (Sigma) for 1hr, washed and then incubated with RMA cells for one hour.

In experiments where PD-L1 was blocked, 5 μg of PD-L1 blocking antibody clone 10F.9G2 (or isotype control) was added to the co-culture.

In experiments where PD-1 was blocked in vitro, tumor cells were incubated with 5 µg of RMP1-14, Fc-silent RMP1-14 or control isotype for 20 minutes, then NK cells were added to the culture. After 2 days, additional 5 µg of antibodies were added to the co-culture and cells were harvested and analyzed after 24 hrs. To check PD-1 saturation, an aliquot of co-culture or tumor cell alone was stained with directly conjugated RMP1-14 or a non-competing PD-1 antibody (29F.1A12).

### Flow cytometry

When needed, tumors were excised from mice, cut in pieces, resuspended in serum-free media, and dissociated using a gentle MACS dissociator (Miltenyi). Following dissociation, the single cell suspension was passed through a 40 µm filter and cells were washed and resuspended in PBS for staining. Spleens were harvested, gently dissociated through a 40 µm filter, washed, and red blood cells were lysed using ACK buffer (Sigma), then washed and resuspended in PBS for staining.

The cellular preparation was stained with the Zombie NIR Fixable Viability Dye (BioLegend) for 20 mins in PBS to label dead cells. Cells were then washed with flow buffer (PBS + 0.5% BSA) and incubated for 20 minutes with purified rat anti-mouse CD16/CD32 (Clone 2.4G2) (BD Biosciences) to block FcγRII/III receptors, followed by washing in flow buffer, and then incubated for a further 20 minutes with primary specific antibodies. Cells were washed and resuspended in flow buffer for sample acquisition or fixed in BD Cytofix/Cytoperm and acquired within 7 days. Flow cytometry was performed using an LSRFortessa (BD) or a Celesta (BD), and data were analyzed with FlowJo software (Tree Star Inc.)

### Antibodies

For experiments with murine cells, the following antibodies were used: *i)* from BD Biosciences: anti-CD3ε (clone 145-2C11); anti-CD8a (clone 53-6.7); anti-CD11b (clone M1/70); anti-CD11c (clone HL3); anti-CD45.2 (clone 104); anti-CD49b (clone DX5); anti-CD69 (clone H1.2F3); anti-Ly6G (clone 1A8); anti-NKR-P1C (clone PK136); anti-Sca-1 (clone D7); *ii)* from Biolegend: anti-CD4 (clone RM4-5); anti-CD19 (clone 6D5); anti-TCRvβ12 (clone MR11-1); anti-Thy-1.1 (clone OX-7); anti-F4/80 (clone BM8); anti-Ly6c (clone HK1.4); anti-NKp46 (clone 29A1.4); anti-PD-1 (clone 29F.1A12); anti-PD-L1 (clone 10F.9G2); rat IgG2a isotype control; and mouse-IgG1 isotype control; *iii)* from Abcam: anti-CD45.1 (clone A20).

For experiments with MM patients, the following antibodies were used: anti-CD3 (clone SK7), anti-CD7 (clone M-T701), anti-CD16 (clone 3G8), anti-CD38 (clone HIT2), anti CD45 (clone HI30), anti-CD56 (clone NCAM16.2), anti-CD138 (clone MI15) and anti-PD1 (clone EH12.1), all from BD Biosciences.

### Generation of cell line variants

RMA, and C1498 cells were transduced with the retroviral expression vector MSCV-IRES-Thy1.1-DEST (Addgene, 17442), by spin infection (800 x g for 2 hours at 37°C) with 8 μg/ml polybrene, and Thy1.1+ cells were sorted.

Single-guide RNA (sgRNA) targeting the first exon of the *Pdcd1* gene (sequence: TGTGGGTCCGGCAGGTACCC) was cloned into the LentiCRISPR lentiviral backbone vector (Addgene 52961), also containing the *Cas9* gene. Lentiviral expression vectors were generated by transfecting 293T cells with 2 μg vector with 2 μg packaging plus polymerase-encoding plasmids using Lipofectamine 2000. Virus-containing supernatants were used to transduce RMA-*Thy1.1* cells by spin infection and PD-1 negative cells were sorted.

C1498-PD-1+ cells were obtained by sorting PD-1+ C1498 parental cells.

Generation of RMA-S-*Pdl1* cells was previously described(*37*).

All engineered cells were regularly assessed for phenotype maintenance by flow cytometry.

### ATAC-seq

Genomic snapshots were generated using IGV software (Broad Institute) using data available on GEO: GSE77695(*53*) and GEO: GSE145299(*54*).

### Analysis of patients

BM aspirates were obtained from patients with clonal plasma cell disorder enrolled at the Ottawa Hospital Research Institute and at the Division of Hematology (“Sapienza” University of Rome). BM samples were lysed using a buffer composed of 1.5 M NH_4_Cl, 100 mM NaHCO_3_, and 10 mM EDTA and then stained as described above.

### Statistical analysis

Differences in tumor growth curves were analyzed with a Two-way ANOVA. Comparison between two groups were performed with Student’s t-test (two tailed, paired or unpaired). Comparison between three groups were performed with ANOVA. p<0.05 was considered a statistically significant difference.

### Study approvals

Mouse studies were reviewed and approved by Animal Care Veterinary Services at the University of Ottawa in accordance with the guidelines of Canadian Institutes of Health Research. For human studies, informed and written consent in accordance with the Declaration of Helsinki was obtained from all patients, and approval was obtained from the Ethics Committee of the Sapienza University of Rome (RIF.CE: 5191) or of The Ottawa Hospital (REB 20180221-02H).

## Results

### Natural Killer cells acquire PD-1 from tumor cells

Consistent with what has been previously been reported (*37, 44*) murine NK cells stimulated ex vivo with a panel of inflammatory mediators failed to upregulate PD-1 at the protein level (Fig. S1). Lack of PD-1 induction was in line with epigenetic analysis of the *Pdcd1* locus, which was not accessible in splenic NK cells, either before or after cytokine stimulation, in sharp contrast with the promoter of another checkpoint receptor (*Tigit*) in NK cells, or *Pdcd1* locus in CD8^+^ T cells (Fig. S2).

Considering the conflicting evidence regarding PD-1 expression on NK cells (*37, 43, 44*) we hypothesized that rather than endogenously expressing the protein, NK cells acquired PD-1 from other cells. To test this hypothesis, we initially used RMA cells, which derive from transformation of murine T cells (*55*), express high levels of PD-1 (Fig. 1A, in red), and were used extensively in our previous study (*37*). We generated RMA cells expressing the syngeneic marker Thy-1.1 (not expressed by C57BL/6 mice, which express the Thy-1.2 allelic variant) and targeted PD-1 with CRISPR/Cas9 (RMA-*Pdcd1^-/-^Thy1.1*) (Fig. 1A, in blue and purple, respectively). We then co-cultured tumor cells with splenocytes from *Pdcd1^+/+^* or *Pdcd1^-/-^* littermates with RMA cells expressing PD-1 or not. In the absence of tumor cells, immune cells did not stain for PD-1 or Thy-1.1. In sharp contrast, NK, T and B cells from both *Pdcd1^+/+^* and *Pdcd1^-/-^* mice stained positively for PD-1 when incubated with RMA cells, but not RMA-*Pdcd1^-/-^* cells (Fig. 1B), indicating that PD-1 was not endogenously expressed by innate and adaptive lymphocytes, but acquired from tumor cells in these settings. Consistent results were obtained by using NK cells isolated from *Pdcd1^+/+^* or *Pdcd1*^-/-^ mice (purity ∼90%) (Fig. 1C and Fig. S3A). Regardless of PD-1 expression on tumor cells, Thy-1.1 was detected in abundance on the surface of immune cells (Fig. 1B-C, Fig. S3B). Acquisition of PD-1 and Thy-1.1 by NK cells was tightly correlated (Fig. S3C), suggesting that the two molecules were transferred to NK cells as part of a unique phenomenon. To determine if other proteins endogenously expressed by RMA cells were acquired by NK cells, we co-cultured CD45.1-expressing NK cells with RMA cells (which express CD45.2). In addition to PD-1 and Thy-1.1, NK cells also acquired TCRvβ12 and CD45.2 (Fig. S4), although the staining was weaker than for PD-1 and Thy-1.1. These data indicate that when interacting with RMA cells, NK cells acquire several proteins they would not endogenously express.

**Figure 1:**
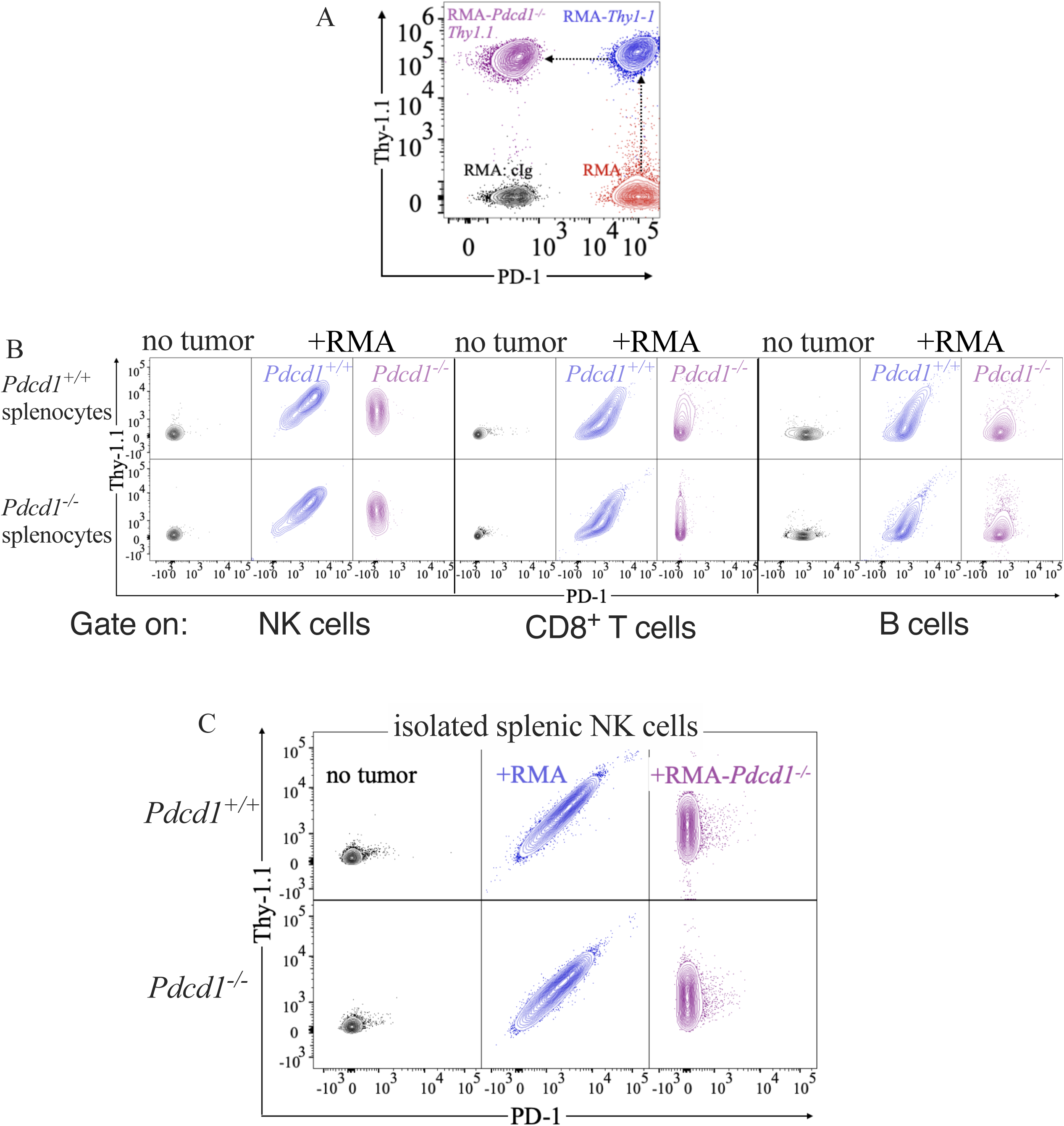
Lymphocytes acquire PD-1 and Thy.1-1 from RMA cancer cells. (A) RMA cells (red) were transduced with a retroviral vector encoding Thy-1.1 to generate RMA-*Thy1.1* (blue) and then PD-1 was knocked-out by CRISPR/Cas9 to generate RMA-*Pdcd1^-/-^Thy1.1* (purple). A representative flow-cytometry staining depicting PD-1 and Thy-1.1 expression is shown. (B) Splenocytes from *Pdcd1^+/+^* or *Pdcd1^-/-^* littermates were incubated with RMA-*Thy1.1* or RMA-*Pdcd1^-/-^Thy1.1*. After 3 days, cells were stained with Thy1.1 and PD-1 antibodies. NK cells were gated as singlets/live-NK1.1^+^NKp46^+^DX5^+^ events. CD8^+^ T cells were gated as singlets/live-CD3^+^CD8^+^ events, B cells as singlets/live-CD19^+^. The experiment depicted is representative of three performed with similar results. (C) NK cells isolated from *Pdcd1^+/+^* or *Pdcd1^-/-^* littermates were incubated with RMA-*Thy1.1* or RMA-*Pdcd1^-/-^Thy1.1*. After 3 days, cells were stained with Thy1.1 and PD-1 antibodies. The experiment depicted is representative of fifteen performed with similar results.

To expand on these results, we next employed C1498 cells, an often-used model of leukemia(*56–58*). A fraction of C1498 cells (∼5%) endogenously expressed PD-1 in culture (Fig. 2A). We sorted PD-1^+^C1498 cells, confirmed that they stably expressed PD-1 upon 2 weeks in culture (Fig. 2B), and then incubated them with splenocytes from *Pdcd1^+/+^* or *Pdcd1^-/-^* littermates. In accordance with the results obtained with RMA cells, both NK cells and CD8^+^ T cells from *Pdcd1^-/-^* mice acquired PD-1 when incubated with C1498 cells, and more so if tumor cells had higher PD-1 expression (Fig. 2C). PD-1 staining observed in *Pdcd1^-/-^* mice was very similar to what observed in the *Pdcd1^+/+^* littermate controls, suggesting that even in the C1498 model, the most PD-1 was not endogenously expressed by immune cells, but rather came from the tumor cells. Similar experiments were repeated using purified NK cells from *Pdcd1^-/-^* NK cells. After 24 hours, NK cells incubated with PD-1+C1498 cells stained positively for PD-1 (Fig. 2D). PD-1 staining was further increased at 72 hours, when we also observed a shift in NK cells incubated with C1498 parental cells (Fig. 2D). Taken together, these data show that NK cells and CD8 T cells acquire PD-1 from leukemia tumor cell lines in vitro.

**Figure 2:**
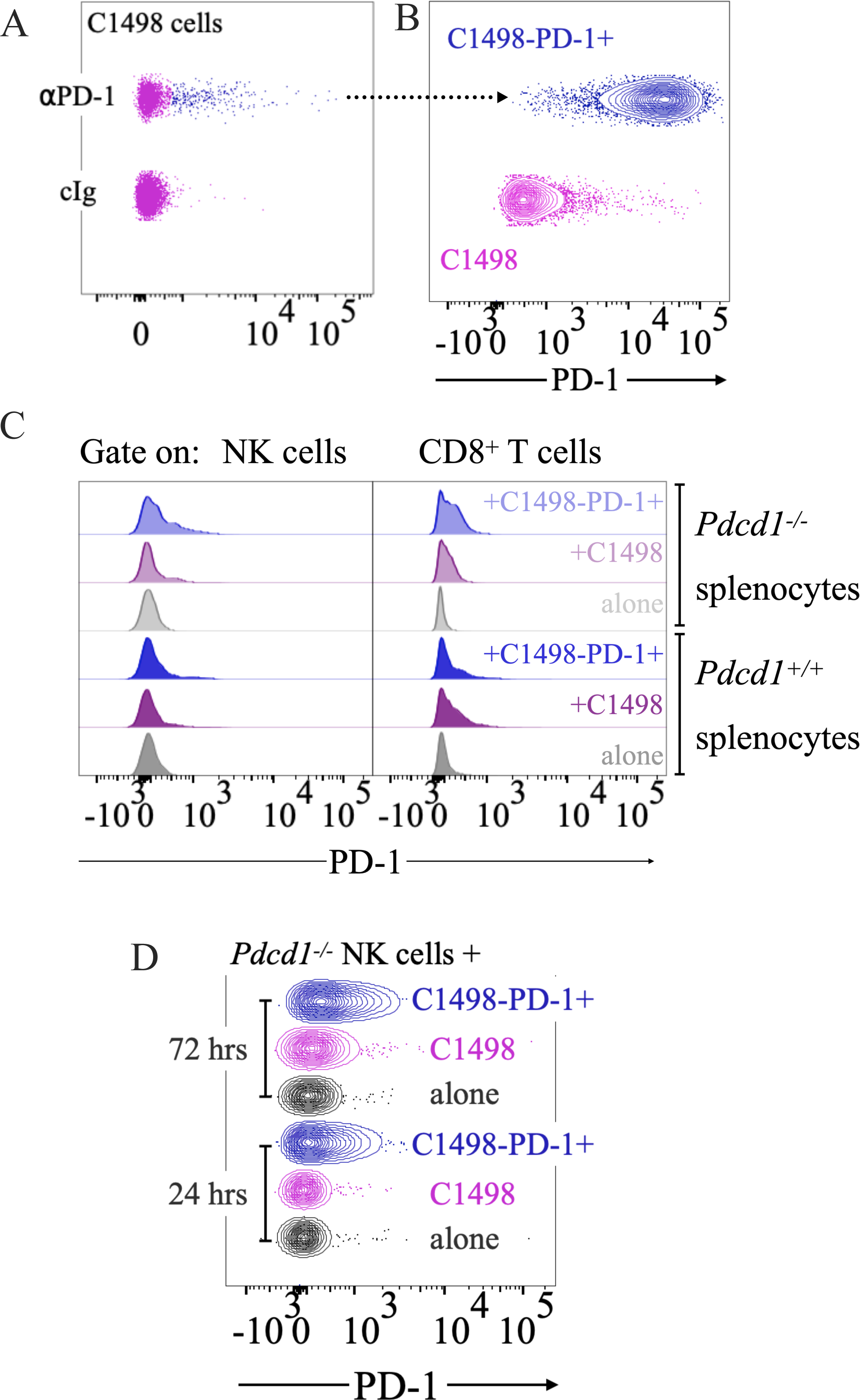
NK cells and T cells acquire PD-1 from C1498 cancer cells. (A) C1498 cells were stained with PD-1 antibody or isotype control. PD-1+ cells (in blue) were flow-sorted and after 2 weeks in culture stained for PD-1, alongside with parental C1498 cells (B). (C) Splenocytes from *Pdcd1^+/+^* or *Pdcd1^-/-^* littermates were cultured with C1498 or C1498-PD-1+ cells for 3 days, and then stained with PD-1 antibodies. NK cells and CD8^+^ T cells were gates as described in 1A. The experiment depicted is representative of three performed with similar results. (D) Splenic NK cells isolated from *Pdcd1^-/-^* mice were co-cultured with C1498 or C1498-PD-1+ cells, or without tumor cells as a control, for 24 hrs or 72 hrs, and stained for PD-1. The experiment depicted is representative of six performed with similar results.

### Trogocytosis is responsible for intercellular transfer of PD-1 from tumor to NK cells

Once we established that NK cells acquired PD-1 from tumor cells, we next investigated whether trogocytosis was responsible for PD-1 transfer. Cell-cell contact is required for trogocytosis. Consistent with our hypothesis that PD-1 is acquired by trogocytosis, *Pdcd1^-/-^* NK cells cultured in transwell with tumor cells (where physical interaction between the two cell types is precluded) failed to stain for PD-1 and Thy-1.1 (Fig. 3A). Further, NK cells incubated with supernatant conditioned by RMA cells failed to stain for PD-1 (Fig. 3B). These experiments reveal that cell-cell contact is required for PD-1 acquisition by NK cells, and suggest that soluble or exosomal PD-1 is not responsible for PD-1 transfer.

**Figure 3:**
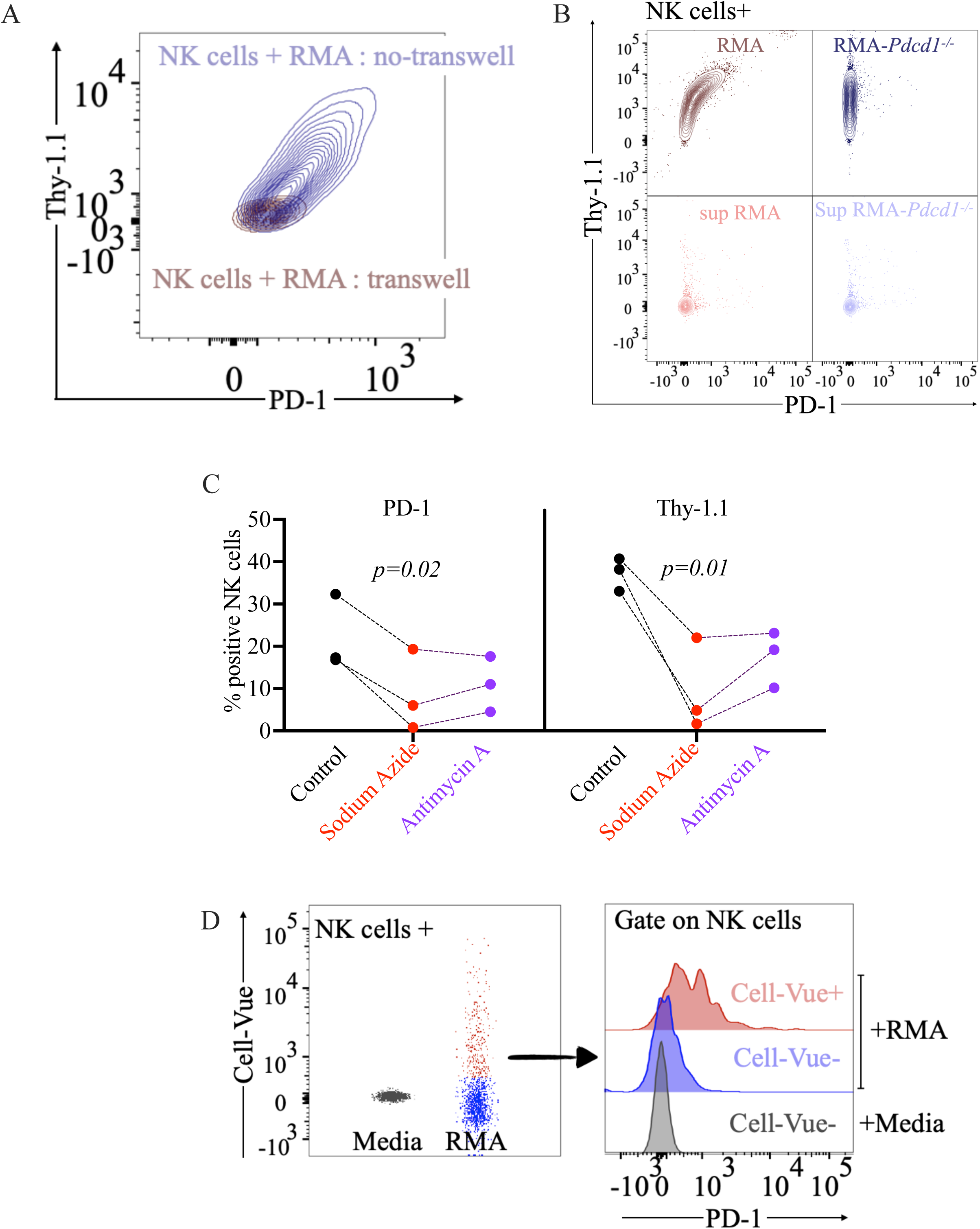
Trogocytosis is responsible for intercellular transfer of PD-1 from tumor to NK cells. (A) Splenic NK cells isolated from a *Pdcd1^-/-^* mouse were co-cultured for 24 hrs with RMA cells separated or not by a transwell semi-permeable membrane before staining for PD-1 and Thy-1.1. The experiment depicted is representative of four performed with similar results. (B) Splenic NK cells isolated from a *Pdcd1^-/-^* mouse were co-cultured for 24 hrs with RMA cells or with media conditioned for three days by RMA cells, then stained for PD-1 and Thy-1.1. The experiment depicted is representative of three performed with similar results. (C) Splenic NK cells isolated from a *Pdcd1^-/-^* mouse were pre-treated with Sodium Azide or Antimycin-A, then co-cultured for one hour with RMA cells and stained for PD-1 and Thy-1.1. Three independent experiments are plotted. Statistical analysis with one-way ANOVA with repeated measurements. (D) NK cells were incubated with RMA cells pre-labelled with Cell-Vue for 24 hrs. Cell-Vue and PD-1 staining on NK cells is depicted, on the left and right respectively.

Blocking ATP synthesis is known to interfere with trogocytosis.(*10*) Consistent with the idea that PD-1 is acquired via trogocytosis by NK cells, pretreatment of NK cells with sodium azide or Antimycin-A, which both prevent ATP synthesis, resulted in a strong reduction of PD-1 and Thy-1.1 acquisition (Fig. 3C).

Transfer of proteins via trogocytosis is accompanied by transfer of membrane lipids. PD-1 transfer was coupled with acquisition of lipids from tumor cells, as revealed by experiments wherein NK cells were co-cultured with RMA cells previously labelled with Cell-Vue, a dye that intercalates in the lipid regions of the cellular membrane (Fig. 3D). Not only did NK cells become robustly positive for the dye, but PD-1 staining was more abundantly detected on NK cells that also acquired lipids from tumor cells (Fig. 3D). These experiments indicate that PD-1, together with other proteins, is acquired contextually with transfer of whole membrane fragments, which is consistent with trogocytosis.

#### SLAM receptors are required for NK cells to trogocytose PD-1 from tumor cells

Acquisition of proteins from donor cells can be facilitated by receptor-ligand engagement, a process known as trans-endocytosis, which NK cells are known to mediate (*59*). In culture, NK cells fail to express PD-L2 but express PD-L1 (Fig. 4A and (*60*)), which could therefore serve as a ligand for trans-endocytosis-driven PD-1 acquisition. However, a saturating dose of PD-L1 blocking antibody did not reduce PD-1 acquisition (Fig. 4B-C) as we would expect if trans-endocytosis was involved. Similar results were obtained blocking PD-1 on RMA cells with an antibody that prevents binding of PD-L1. Despite the antibodies saturated PD-1 on the membrane of RMA cells (Fig. S5A), PD-1 and Thy-1.1 were still effectively transferred to NK cells (Fig. S5B). These experiments not only suggest that PD-1/PD-L1 binding is not required for PD-1 transfer, but also imply that Fc-receptor engagement by PD-1 antibodies does not facilitate trogocytosis (*61*). Finally, we sought genetic corroboration using PD-L1-deficient NK cells from two different mouse strains: a full body PD-L1 knock-out (*Cd274^-/-^*) (*48*) and an NK cell specific PD-L1 knockout (*Ncr1^+/Cre^* x *Cd274^fl/fl^*) (*47*) that we crossed with PD-1-deficient mice (*Pdcd1^-/-^ Ncr1^+/Cre^ Cd274^fl/fl^*). PD-L1 deficient NK cells acquired PD-1 and Thy-1.1 at levels similar to NK cells isolated from PD-L1 expressing controls (Fig. 4D-E). PD-L1 was also dispensable for PD-1 and Thy-1.1 acquisition in CD8^+^ T cells and B cells (Fig. S6).

**Figure 4:**
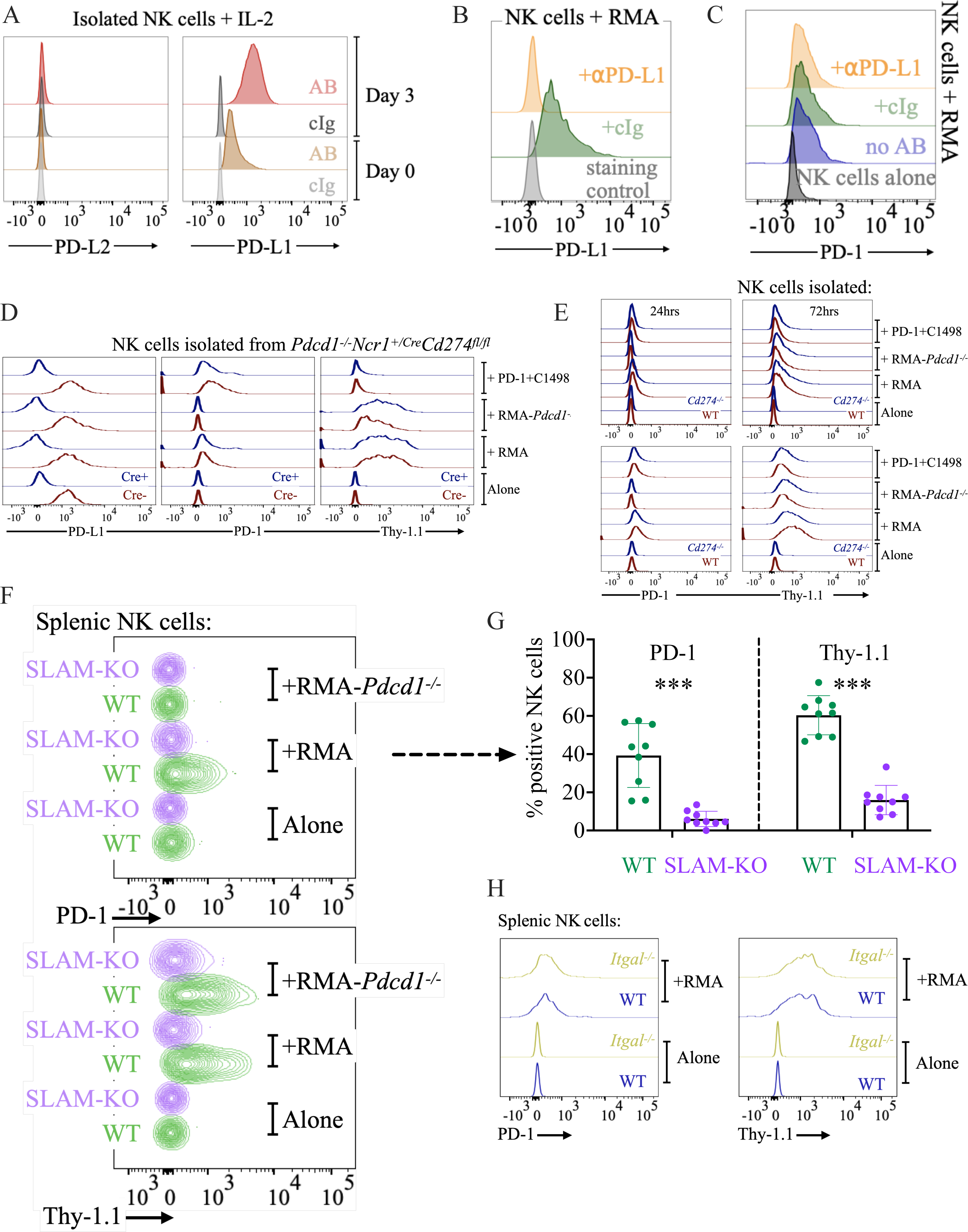
SLAM receptors are essential for trogocytosis. (A) Splenic NK cells isolated from a *Pdcd1^-/-^* mouse were cultured for 3 days and then PD-L2 and PD-L1 expression was analyzed by flow-cytometry. Representative of three experiments performed with similar results. (B-C) NK cells were incubated with RMA cells in the presence of a PD-L1 blocking antibody or an isotype control for 24 hrs, before being stained for PD-1 and PD-L1. As additional controls, NK cells were: *i)* co-cultured with RMA without adding any antibody; or *ii)* cultured alone without adding tumor cells. The experiment depicted is representative of three performed. (D-E) NK cells were isolated from the spleen of *Pdcd1^-/-^Ncr1^+/Cre^Cd274^fl/fl^* (D) or *Cd274^-/-^* (E) mice and co-cultured for three days with RMA or C1498 tumor cells, when PD-1 and Thy-1.1 staining was assessed by flow cytometry. (F-G) Splenocytes from SLAM-deficient mice or control littermates were co-cultured for three days with RMA or C1498 cells. PD-1 and Thy-1.1 staining on NK cells was then assessed by flow cytometry. The experiment depicted is representative of 3 performed. In G, the frequency of NK cells staining for PD-1 or Thy-1.1 in WT or SLAM-deficient mice analyzed in the three experiments is plotted. Statistical analysis with two-tailed unpaired Student’s t-test. ***: p<0.001. (H) Splenocytes from LFA-1-deficient or WT mice were co-cultured for three days with RMA cells. Representative of 3 experiments performed.

SLAM receptors are important mediators of cell-cell interactions between hematopoietic cells and are abundantly expressed not only by NK cells but also by T and B cells (*62*). Given the broad expression of SLAM family members, their importance in regulating the activation of different immune cells and considering that PD-1 was trogocytosed by both innate and adaptive lymphocytes, we hypothesized that SLAM receptors promoted PD-1 trogocytosis. To test this hypothesis, we cultured splenocytes from mice where the whole *SLAM* locus was deleted (*49*) with tumor cells and then assessed PD-1 and Thy-1.1 staining on NK, T and B cells. Consistent with our hypothesis, NK cells from SLAM-deficient mice failed to acquire PD-1 from RMA cells (Fig. 4F-G). Not only was PD-1 acquisition abolished, but more broadly, SLAM-deficient NK cells failed to perform trogocytosis with tumor cells, as revealed by lack of Thy-1.1 transfer (Fig. 4F-G). In addition to NK cells, SLAM-deficient T and B cells also displayed reduced trogocytosis (Fig. S7), confirming that SLAM receptors are key mediators for trogocytosis between immune cells and leukemia cells.

Given the importance of SLAM receptors in mediating cell-cell interactions, we analyzed if deficiency in other adhesion molecules also interfered with trogocytosis. LFA-1 is a key adhesion molecule, but NK cells lacking expression of CD11a,(*50*) a subunit of LFA-1, did not present a deficit in PD-1 or Thy-1.1 acquisition (Fig. 4H). Similar results were also observed analyzing CD8^+^ T and B cells (Fig. S8). NKG2D, an activating receptor ubiquitously expressed by NK cells,(*63*) was also not involved in mediating trogocytosis between NK and RMA cells (Fig. S9). Taken together, these results indicate that SLAM receptors, but not other adhesion molecules or activating receptors, mediate PD-1 trogocytosis from tumor to immune cells.

### Activated NK cells acquire PD-1 from tumor cells in vivo

Next, we performed in vivo studies to determine if intratumoral NK cells trogocytosed PD-1. We injected *Pdcd1^+/+^* or *Pdcd1^-/-^* littermates with RMA or RMA-*Pdcd1^-/-^* cells, both expressing Thy-1.1, and when tumors reached ∼300 mm^3^ we analyzed PD-1 staining on intratumoral lymphocytes. In all cohorts of mice, NK cells infiltrating the tumors stained intensely for Thy-1.1 (Fig. 5A-D, Y axis), showing that trogocytosis occurred in vivo. Strikingly, high levels of PD-1 were detected on the surface of NK cells only when tumor cells expressed PD-1, not only in *Pdcd1^+/+^*, but also in *Pdcd1^-/-^* mice (Fig. 5A and C vs B and D). These data show not only that PD-1 is acquired by tumor infiltrating NK cells, but also that trogocytosis is the major mechanism leading to PD-1 presence on the surface of NK cells in the RMA model. Consistent with what was observed on NK cells, CD8^+^ T cells from *Pdcd1^-/-^* mice also acquired Thy-1.1 and PD-1 from tumor cells (Fig. 5C). As expected, PD-1 staining in CD8^+^ T cells was also observed in *Pdcd1^+/+^* mice injected with PD-1-deficient RMA cells, confirming that CD8^+^ T cells endogenously expressed PD-1 in vivo. In our previous study, we reported that PD-1 staining was higher on activated NK cells (*37*). Analysis of NK and T cells from *Pdcd1^-/-^* mice infiltrating RMA tumors confirmed that PD-1^+^ NK and T cells also stained more brightly for activation markers such as Sca-1 and CD69 (Fig. 5E), supporting the idea that the NK cells that are activated by the encounter with tumor cells are also the ones more susceptible to acquiring PD-1 and therefore being inhibited by it.

**Figure 5:**
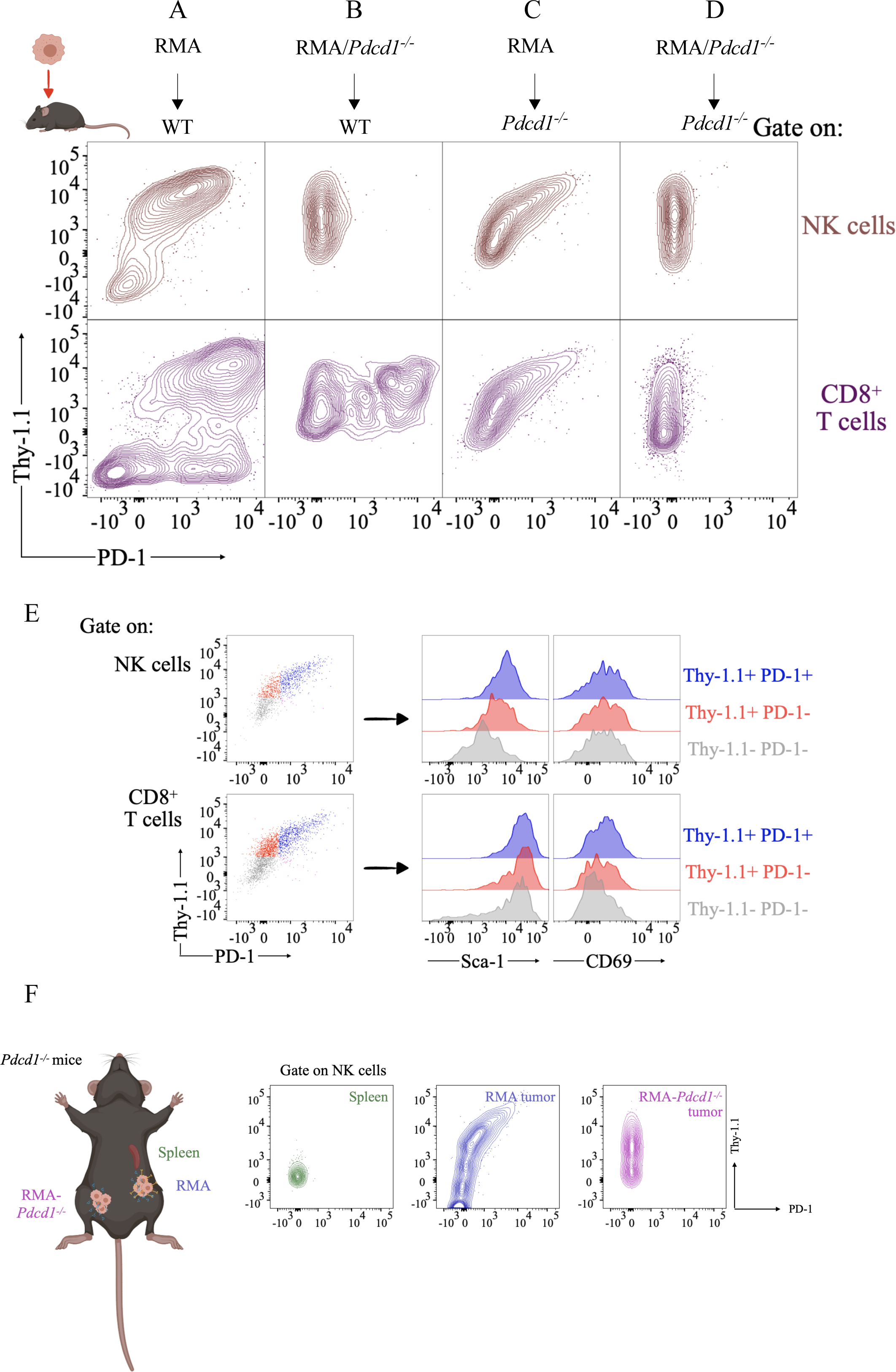
**Intratumoral lymphocytes acquire PD-1 from tumor cells**. (A-D) *Pdcd1^+/+^* or *Pdcd1^-/-^* mice were injected with RMA or RMA-*Pdcd1^-/-^* tumors. PD-1 and Thy-1.1 staining was assessed by flow-cytometry on tumor infiltrating NK and T cells (gated as in Fig. 1B and 1C, respectively). The experiment shown is representative of four performed with similar results. (E) Expression of Sca1 and CD69 was analyzed on *Pdcd1^-/-^* NK and T cells infiltrating RMA tumors, by gating on Thy-1.1^-^PD-1^-^ (gray), Thy-1.1^+^PD-1^-^ (red) or Thy-1.1^+^PD-1^+^ (blue) cells. The mouse depicted is the same depicted in 2C. (F) RMA or RMA-Thy.1-1 cells were injected in either flank of a *Pdcd1^-/-^* mouse. PD-1 and Thy.1-1 staining was analyzed in intratumoral or splenic NK cells. The experiment depicted is representative of two performed.

To further support our hypothesis that PD-1 was trogocytosed by NK cells in the tumor microenvironment, we injected *Pdcd1^-/-^* mice with RMA or RMA-*Pdcd1^-/-^* cells in either flank. As previously reported (*37*), splenic NK cells failed to stain for PD-1 (Fig. 5F). Consistent with what is described in Fig. 5A-D, NK cells in both tumors acquired Thy-1.1 from tumor cells, but only NK cells in PD-1 expressing tumors also stained for PD-1 (Fig. 5F). Taken together, these results highlight how activated NK cells perform trogocytosis and acquire PD-1 in the tumor microenvironment in vivo.

### PD-1 acquired via trogocytosis inhibits NK cell responses against cancer

Once we established that NK cells trogocytose PD-1 in vivo, we sought to determine if trogocytosed PD-1 suppressed anti-tumor immunity. For these studies, rather than using RMA cells (which are resistant to both T and NK cell responses (*37, 55, 56, 64*)) we took advantage of RMA-S-*Pdl1* cells we previously generated (*37*). Similar to RMA, these cells express high levels of PD-1, but differently than RMA, they lack MHC class I expression and are therefore susceptible to NK-mediated control (*37*). Using CRISPR/Cas9, we generated an RMA-S-*Pdl1* variant lacking PD-1 expression (Fig. 6A). When injected in *Pdcd1^-/-^* mice, where the only source of PD-1 on NK cells are tumor cells, we observed a dramatic deceleration in outgrowth of tumor cells lacking PD-1 expression (Fig. 6B). However, lack of PD-1 did not delay cell growth in vitro, nor it prevented RMA-S-*Pdl1-Pdcd1^-/-^* cells from growing tumors in immunodeficient mice (Fig. 6C). These data indicate that, rather than having cell intrinsic growth defects, PD-1-deficient tumor cells have reduced capacity of forming ectopic tumors as they fail to inhibit NK cells via PD-1 transfer.

**Figure 6:**
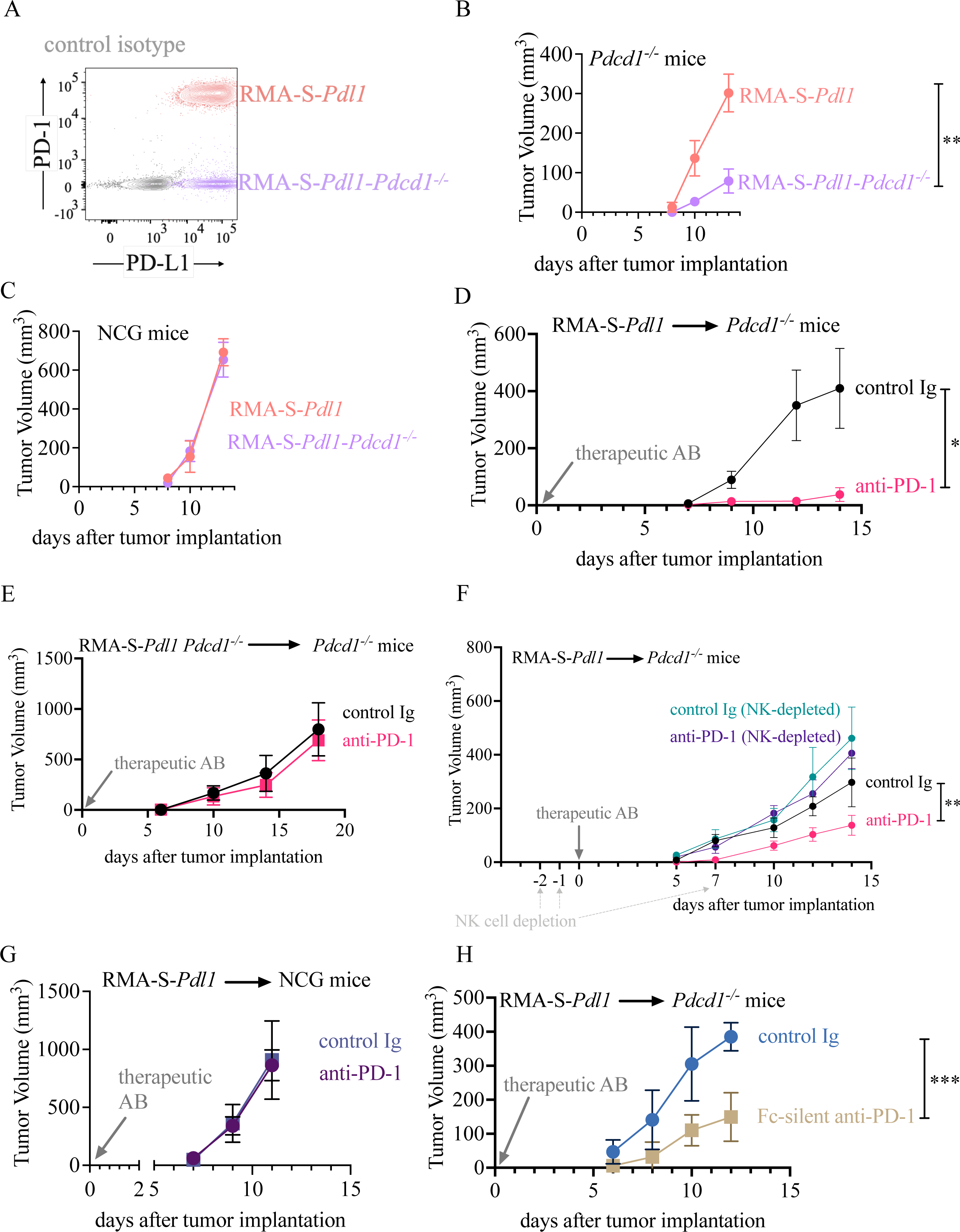
PD-1 blockade is effective in *Pdcd1^-/-^* mice when NK cells are present and tumor cells express PD-1. (A) PD-1 and PD-L1 expression in RMA-S-*Pdl1* or RMA-S-*Pdl1-Pdcd1^-/-^* cells was analyzed by flow cytometry. (B-H) In all experiments, the indicated cell lines were injected resuspended in Matrigel, alone or mixed with different PD-1 blocking or control antibody. Tumor growth was assessed over time and data were analyzed with 2-way ANOVA. (B) n=6/group, the experiment depicted is representative of two performed with similar results. (C) n=6/group, the experiment depicted is representative of two performed with similar results. (D) n=4/group, the experiment depicted is representative of two performed with similar results. (E) n=6/group, the experiment depicted is representative of three performed with similar results. (F) n=at least 5/group, the experiment depicted is representative of two performed with similar results. (G) n= 5/group, the experiment depicted is representative of two performed with similar results. (H) n= 5/group, the experiment depicted is representative of two performed with similar results. *: p<0.05; **:p<0.01; ***:p<0.001.

Using the RMA-S-*Pdl1* model, we previously showed that PD-1 blockade rescued the ability of NK cells to control tumor growth in vivo (*37*). Considering that PD-1 expression in tumor cells promoted in vivo growth in a cell extrinsic fashion, we reasoned that the therapeutic effect of PD-1 blockade should also be observed in *Pdcd1^-/-^* mice. In accordance with our hypothesis, when we treated PD-1-deficient mice injected with RMA-S-*Pdl1* cells with a PD-1 blocking antibody (RMP1-14) we observed a dramatic reduction in tumor outgrowth (Fig. 6D). On the other hand, when we injected PD-1-deficient mice with PD-1 deficient RMA-S-*Pdl1* cells, PD-1 blockade had no therapeutic effect (Fig. 6E).

To confirm that NK cells, and not other components of the immune response, were inhibited by tumor derived PD-1, we injected RMA-S-*Pdl1* cells in mice where NK cells were depleted using a monoclonal antibody (PK136) and treated the mice with PD-1 blockade. NK cell depletion was sufficient to abolish the therapeutic effect of the blocking antibody, whereas PD-1 blockade delayed tumor outgrowth in the control group (Fig. 6F). Corroboration of these results came from experiments where PD-1 antibodies failed in immunocompromised mice (Fig. 6G). Moreover, the similar in vivo growth of tumor cells in immunocompromised mice (Fig. 6G) excluded a tumor cell intrinsic effect of PD-1 blocking antibodies.

Finally, to rule out that the therapeutic effect of PD-1 antibodies was due to Antibody-Dependent Cellular Cytotoxicity (ADCC) potentially mediated by NK cells against cancer cells coated with PD-1 antibodies, we employed an engineered version of anti-PD-1 that lacks the ability to bind to Fc-receptors (Fc-silent RMP1-14) (*52*). Treatment with Fc-silent PD-1 antibodies delayed the outgrowth of PD-1 expressing tumors (Fig. 6H), indicating that the therapeutic effect of PD-1 antibodies in *Pdcd1^-/-^* mice was not due to ADCC. Taken together, these results indicate that trogocytosed PD-1 inhibits the anti-tumor activity of NK cells, which can be rescued by PD-1 blocking antibodies.

### Identification of an NK cells population in multiple myeloma patients staining for tumor cell markers and PD-1

Finally, to determine if NK cells acquire PD-1 from tumor cells in cancer patients, we analyzed PD-1 staining in NK cells from the bone marrow (BM) of patients with clonal plasma cell disorders (Table 1 details patients’ information). Pathological analysis showed that of the 28 patients analyzed, 21 were diagnosed with multiple myeloma (MM), 3 with monoclonal gammopathy of undetermined significance (MGUS), 3 with smoldering myeloma, and 1 with solitary plasmacytoma. For these studies, we relied on CD138, a protein frequently expressed by clonal plasma cells but not by NK cells (*65*), as a surrogate trogocytosis marker. Flow cytometry analysis confirmed the presence in several patients of a high scattering CD138^+^ population (Fig. S10), that we identified as clonal plasma cells. In support of our gating strategy, patients diagnosed with MM presented a higher CD138^+^ frequency than MGUS patients. NK cells were instead gated as low scattering events, live, CD3^-^CD56^+^CD16^+/-^ and in most samples CD7 was also used (Fig. S10). In the vast majority of samples analyzed, high scattering-CD138^+^ cells expressed PD-1 (Sup. Fig. 10). Indication that NK cells performed trogocytosis came from analysis of BM aspirates where a CD138^+^ NK cell population, in both CD56^bright^CD16^-^ and CD56^dim^CD16^+^ NK cell subsets, was identified (Fig. 7, in pink and green), corroborating the results obtained in murine models. Notably, we found a sizeable and consistent (albeit often small) population of NK cells that stained for both CD138 and PD-1 (Fig. 7, in green), supporting the idea that NK cells in patients with clonal plasma cell disorders acquire PD-1 and cancer cell markers from tumor cells.

**Figure 7:**
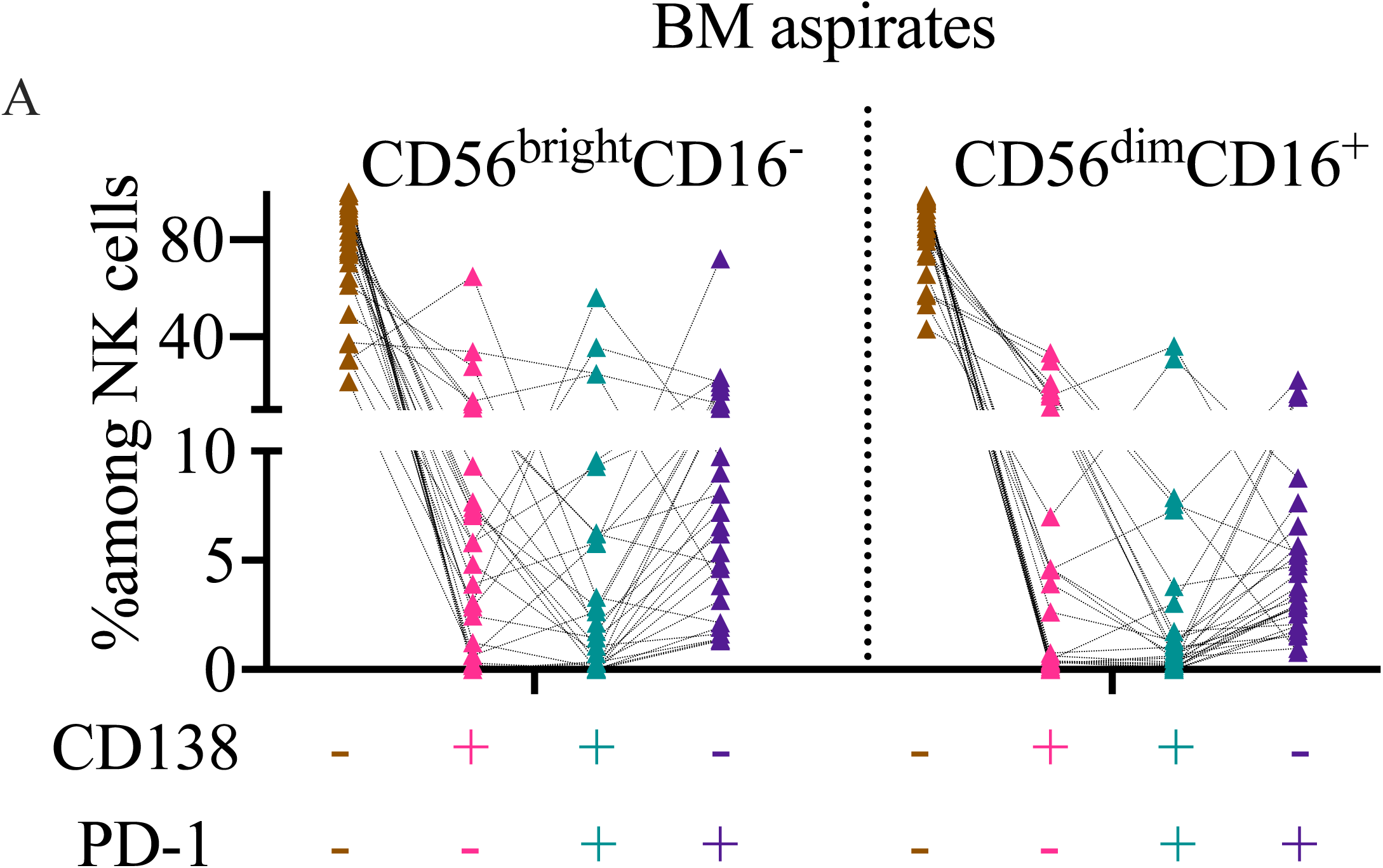
NK cells co-stain for CD138 and PD-1 in the bone marrow of patients with clonal plasma cell disorders. The bone marrow aspirates of 28 patients with clonal plasma cell disorders were analyzed by flow cytometry. The frequency of NK cells staining for either, neither or both CD138 and PD-1 is depicted.

**Table 1.**
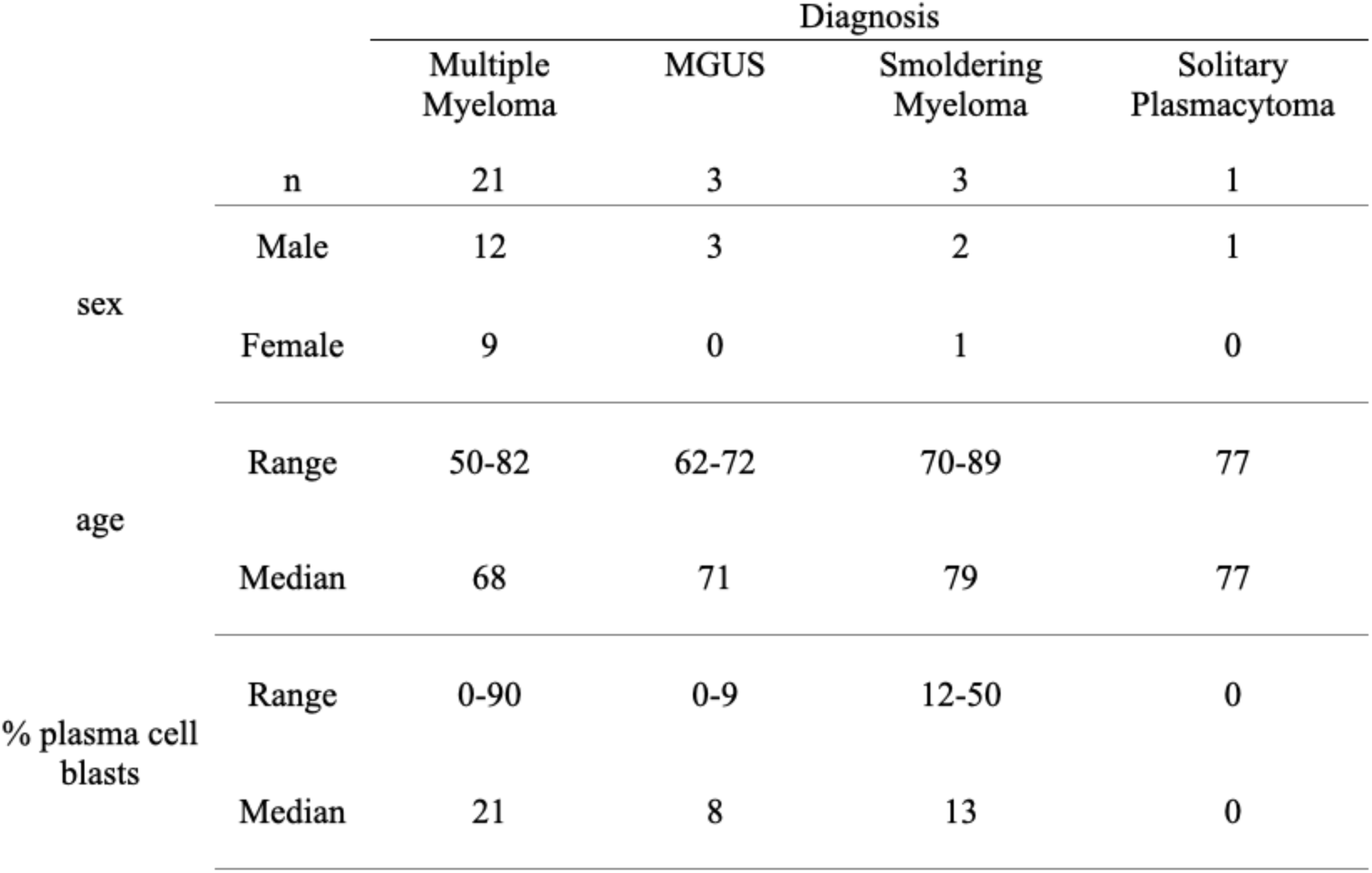

In conclusion, this study identifies trogocytosis as a new mechanism by which PD-1 is acquired from tumor cells. by NK and T cells. PD-1 trogocytosis strongly relies on SLAM receptors and functionally suppresses the ability of NK cells to eliminate tumors in vivo.

## Discussion

The nature of PD-1 expression on NK cells remains fairly elusive, with contrasting evidence indicating that PD-1 is either expressed on not expressed in NK cells (*43, 44*). Considering the importance of PD-1 in suppressing the immune response to cancer (*66*), and given the tremendous interest in the development of NK cell based cancer immunotherapies (*32, 67*), understanding whether PD-1 directly inhibits NK cell function is of the utmost importance. We recently reported that PD-1 suppresses NK cells in several mouse models of cancer (*37*), but previously have not yet deciphered the mechanisms leading to PD-1 upregulation in murine NK cells infiltrating lymphoma mouse models. The lack of PD-1 induction in NK cells following several ex vivo stimulations, combined with the analysis of the *Pdcd1* locus in resting and cytokine stimulated NK cells, prompted us to hypothesize that PD-1 was not endogenously expressed by NK cells but rather be derived from other sources. Several cellular processes have been shown to be responsible for protein transfer. Amongst these processes, trogocytosis, the intercellular exchange of whole membrane fragments, is highly performed by NK cells (*10–12*). Proteins transferred via trogocytosis can have a substantial impact in immune function (*1, 2*). In cancer, trogocytosis has been associated with reduced immune responses and with the failure of immunotherapy. For example, a recent study highlighted how CAR T cells trogocytose antigens from tumor cells and become susceptible to fratricide, greatly limiting the response to cellular therapy (*68*). Trogocytosis triggered by Fc-receptors engaging therapeutic antibodies, performed by myeloid and NK cells, has been a major hurdle limiting the efficacy of monoclonal antibodies against cancer antigens (*15, 61, 69-85*). Despite such evidence and the immense interest in elucidating mechanisms underlying resistance to PD-1 blockade, whether PD-1 is trogocytosed by immune cells has been largely unexplored. Here we show that in some contexts trogocytosis is a major mechanism by which PD-1 becomes localized on the surface of immune cells. This was true not only for NK cells, but also for adaptive lymphocytes. PD-1 acquisition happened in a cell-cell contact dependent fashion, contextualized within the transfer of other proteins and whole membrane fragments and was strongly suppressed by ATP depletion, indicating that PD-1 was trogocytosed by immune cells. Interestingly, PD-1 antibodies did not elicit PD-1 trogocytosis by NK cells, suggesting that PD-1 could be acquired by NK cells even in the absence of Fc-receptor engagement. Mechanistic studies using blocking antibodies and transgenic mice allowed us to exclude a role for PD-L1, abundantly expressed by NK cells, in PD-1 acquisition, ruling out trans-endocytosis as a mechanism of PD-1 transfer. On the other hand, receptors belonging to the SLAM family proved to be essential for intercellular transfer of PD-1 from tumor to immune cells. SLAM receptors are important regulators of immune function and ubiquitously expressed by NK cells (*62*), but also by tumors of hematopoietic origin, including multiple myeloma (*86*). Our finding that SLAMs promote the transfer of PD-1 from tumor to immune cells requires consideration of trogocytosis as an important biological variable when designing mono -or combination therapies targeting these receptors.

Trogocytosed PD-1 was functional and suppressed the anti-cancer activity of NK cells. The in vivo studies performed here further expand on our previous findings that NK cells contribute to the therapeutic efficacy of PD-1 blockade (*37*), and explain why checkpoint blockade relies on NK cells despite their lack of PD-1 expression.

While more translational studies are required to follow up on this mechanistic data, we successfully identified a subset of NK cells which stained for CD138 in the bone marrow of patients with clonal plasma cell disorders. As CD138 is not expressed by NK cells, we relied on CD138 staining to identify bone marrow NK cells that performed trogocytosis. Consistent with our in vivo results, CD138^+^ NK cells also stained for PD-1, and flow cytometry and bioinformatic analysis of a published dataset indicated that multiple myeloma cells can express PD-1 (*87*). Based on our in vivo results, we propose that PD-1 expression, in addition to benefiting cancer cells with intrinsic signaling (*88*), also promotes immune escape. In fact, tumor cells expressing PD-1 can donate this powerful inhibitory receptor to activated immune cells when they are in direct contact. PD-1 acquisition can however be therapeutically abrogated by checkpoint blockade, potentially rescuing the ability of NK cells to promote anti-cancer immunity.

In our current analysis (and differing from our murine results) PD-1 was also found in a fraction of human NK cells that did not stain for CD138. These data are consistent with the idea that human NK cells endogenously express PD-1, as recently corroborated in healthy donors and patients undergoing hematopoietic stem cell transplantation (*43*). Endogenous expression of PD-1 does not exclude the possibility that immune cells also rely on trogocytosis to gain further PD-1 protein from neighbor cells. This notion is well supported by a recent study that identified trogocytosing NK cells in a broad spectrum of hematopoietic malignancies (*89*). In accordance with our data, NK cells labelled with tumor cell markers also stained for PD-1 (*89*). Whether endogenously expressed by NK cells or acquired from cancer or other immune cells, several reports, including the present one, have highlighted the importance of PD-1 in suppressing NK cells (*37–43*).

Finally, in light of these results, it will be important for future immune-profiling efforts based on transcriptomic analysis to take into account that proteins are acquired, sometimes at surprisingly high levels, by immune cells in the tumor microenvironment. Pursuant to our previous studies and given its known importance in suppressing anti-cancer responses we focused on PD-1; however, it is conceivable that other proteins with immunomodulatory potential will be acquired by NK and T cells while interacting with tumor cells. Further characterization of the mechanisms underlying membrane transfer and identification of molecules transferred to immune cells is required to elucidate how immune cells are regulated by checkpoint receptors, and other proteins, in a transcription-independent fashion.

## Acknowledgment

We thank members of the Ardolino lab, Drs. Horan, Vanderhyden, Gray, Scott, Bourgeois-Daigneault, Kissov and Piconese for critically reading the manuscript; Fernando Ortiz, Drs. Tang, Stanford, Ito and the CHEO flow-core for support with flow-cytometry; the ACVS facility at the University of Ottawa for support with animal studies. MA is supported by Ride for Dad, CIHR and Myeloma Canada, HYS by the NIH intramural research programs, GS and AS by AIRC, AZ by Sapienza University, PGF by Science Foundation Ireland, AHS by NIH (PO1 grant 56299). EV is supported by an AIRC fellowship. MM is the recipient of a CAAIF postdoctoral fellowship. JJH and DC are recipients of a Frederick Banting and Charles Best Canada Graduate Scholarships Doctoral Award from CIHR. We are thankful to Dr. Ravetch (Rockefeller University) for providing us with Fc-silent PD-1 antibodies and to Dr. Raulet, for providing us with mouse lines and continuous support. This manuscript is dedicated to the memory of Dr. Lucas Horan, who loved science and the Bay.

## Authorship contributions

Author contributions are detailed according to CRediT criteria.

| Author | Conceptualization | Formal analysis | Funding acquisition | Investigation | Methodology | Resources | Supervision | Visualization | Writing - original draft | Writing review - & editing |
| --- | --- | --- | --- | --- | --- | --- | --- | --- | --- | --- |
| MSH | X |  |  | X | X |  |  |  |  | X |
| MM |  | X |  | X |  |  |  |  |  | X |
| JJH |  |  |  | X |  |  |  |  |  | X |
| EV |  |  |  | X |  |  |  |  | X | X |
| HYS | X | X | X |  |  |  |  | X |  | X |
| AS |  |  |  | X |  |  |  |  |  | X |
| OMM |  |  |  | X |  |  |  |  |  | X |
| FGA |  |  |  | X |  |  |  |  |  | X |
| KPB |  |  |  |  |  | X |  |  |  | X |
| DC |  | X |  |  |  |  |  |  |  | X |
| MTP |  |  |  |  |  | X |  |  |  | X |
| AS |  |  | X |  |  | X | X |  |  | X |
| PGF |  |  |  |  |  | X |  |  |  | X |
| AS |  |  |  |  |  | X |  |  |  | X |
| GS | X | X | X |  |  |  | X |  |  | X |
| AV |  |  |  |  |  | X |  |  |  | X |
| AZ | X |  | X |  |  | X | X |  |  | X |
| AMC | X |  | X |  |  | X |  |  |  | X |
| MA | X | X | X | X | X |  | X | X | X | X |

## SUPPLEMENTARY MATERIAL

**Supplementary Figure 1:**
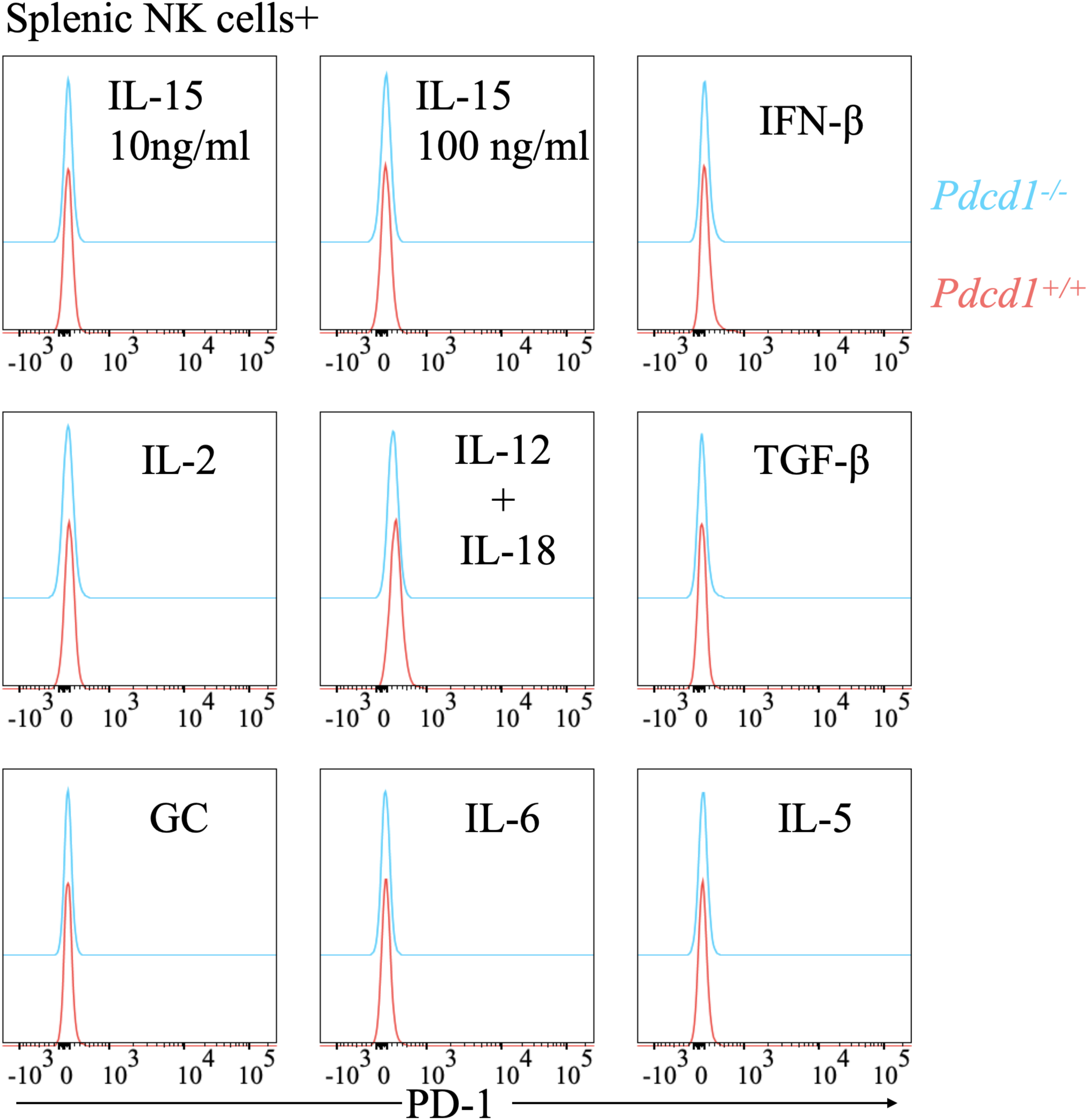
Inflammatory signals fail to induce PD-1 upregulation in purified NK cells in vitro. Magnetically enriched NK cells from *Pdcd1^+/+^* or *Pdcd1^-/-^* mice were cultured for 72hrs with the inflammatory mediators indicated in the figure. GC=glucocorticoid (Corticosterone). NK cells from 3 mice/genotype where pooled. The experiment depicted is representative of 3 performed.

**Supplementary Figure 2:**
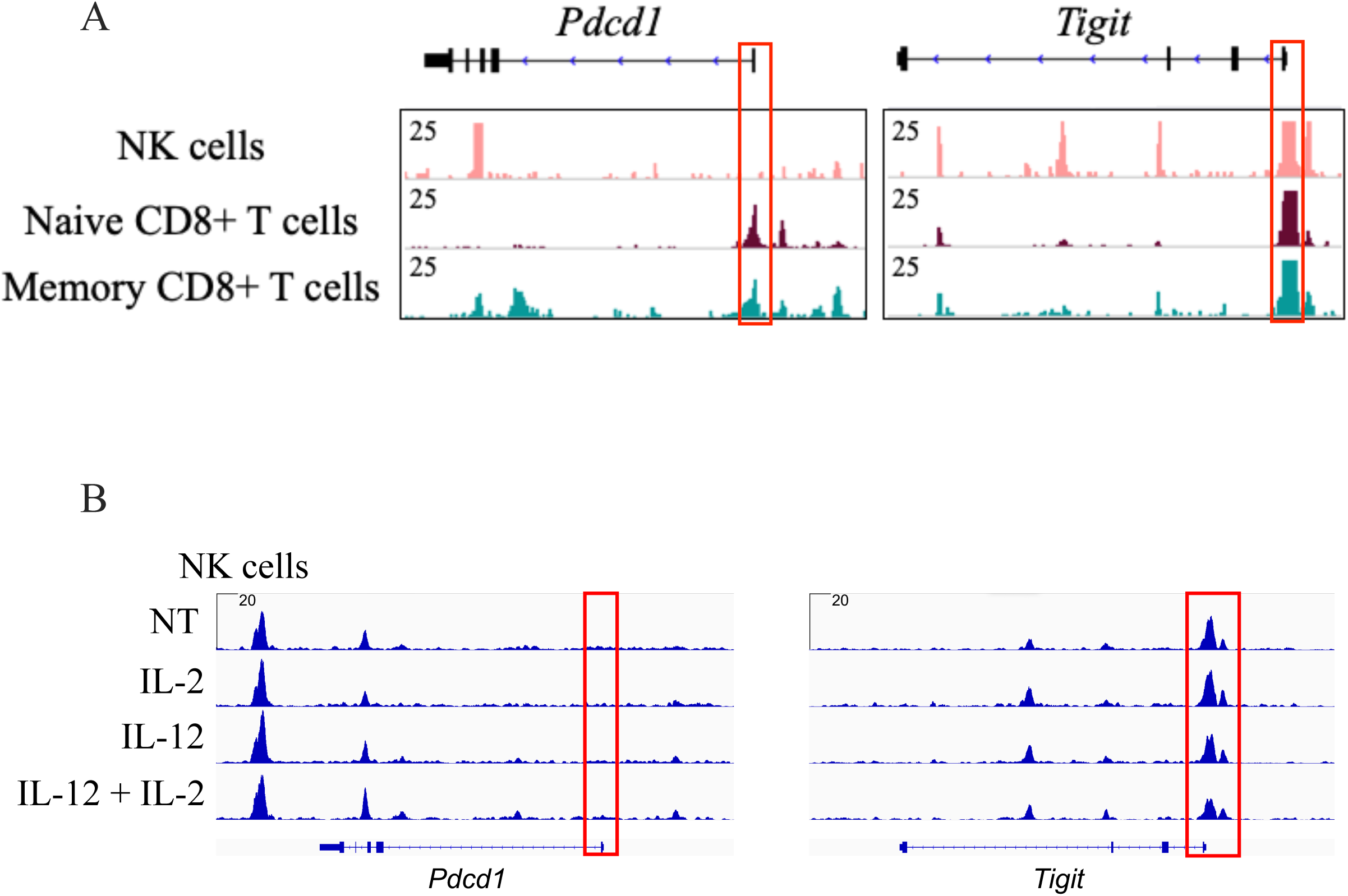
The *Pdcd1* locus is closed in resting NK cells. Genomic snapshots of normalized ATAC-seq signals in NK cells, naïve and memory CD8+ T cells across *Pdcd1* and *Tigit* loci.

**Supplementary Figure 3:**
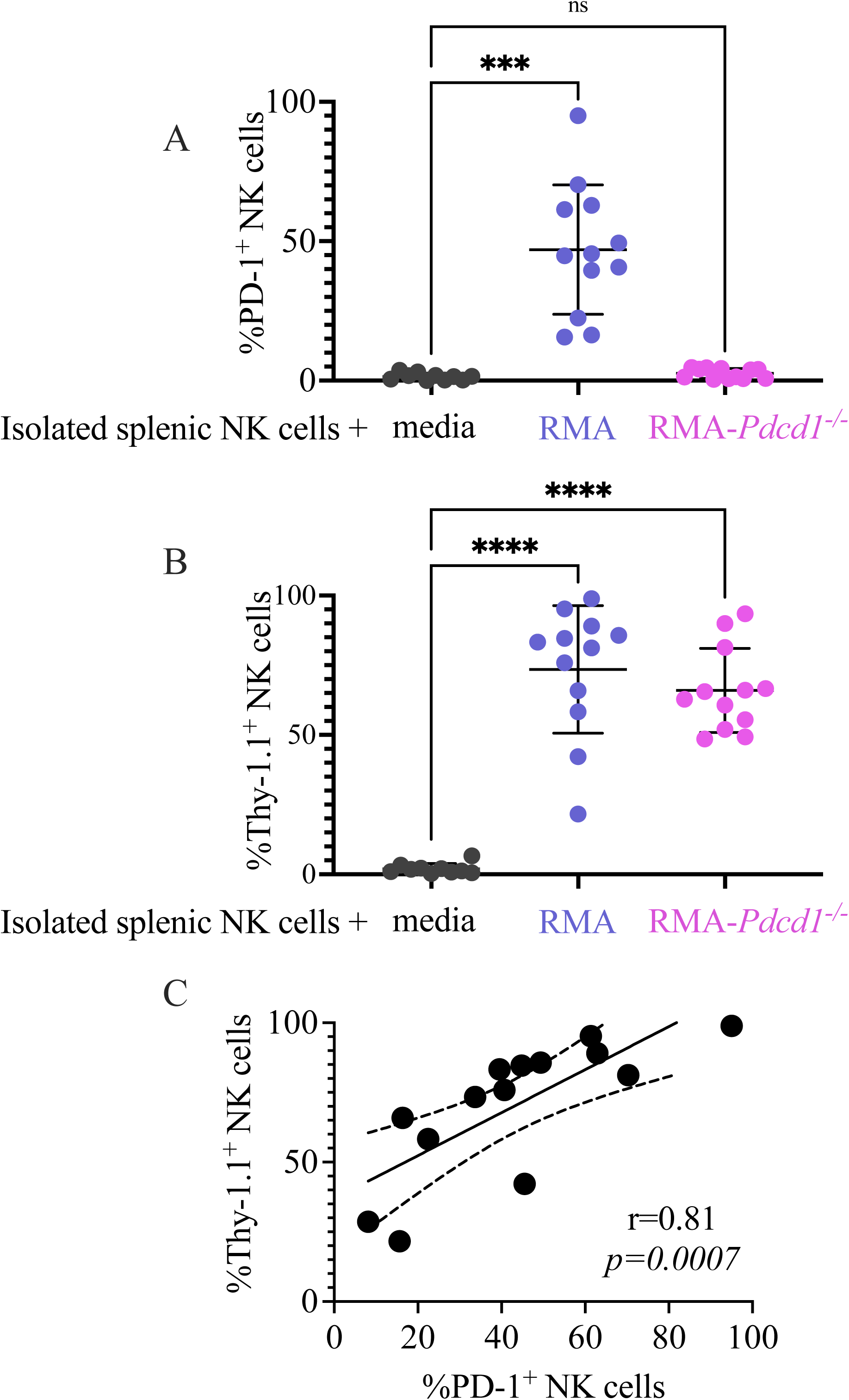
PD-1 and Thy-1.1 are co-acquired by NK cells. NK cells isolated from the spleens of *Pdcd1^-/-^* mice were co-cultured with RMA or RMA-*Pdcd1^-/-^* cells for three days. A and B show the frequency of PD-1^+^ or Thy-1.1^+^ NK cells in the 14 mice analyzed in the 13 experiments performed. In A and B Statistical analysis with ANOVA with Dunnet’s multiple comparison test. In (C) the correlation between PD-1^+^ and Thy-1.1^+^ NK cells is depicted. 95%-confidence interval is also showed, statistical analysis with Spearman correlation test.

**Supplementary Figure 4:**
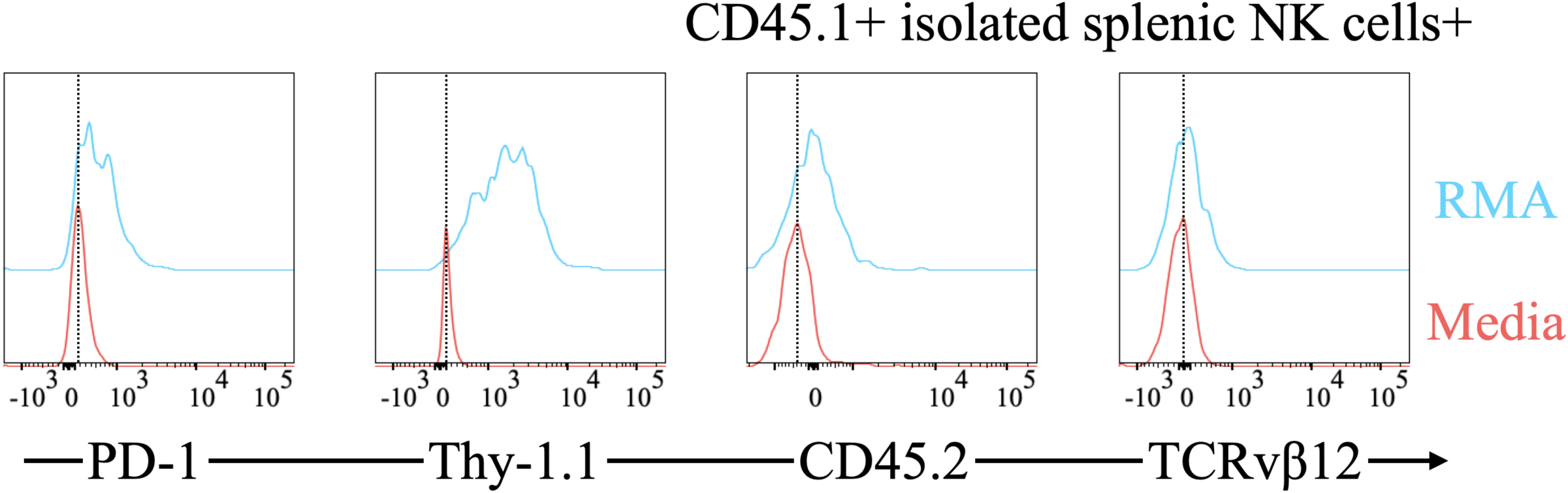
NK cells acquire at least four proteins they do not endogenously express from RMA cells. CD45.1+ NK cells were co-cultured with RMA cells for three days and then PD-1, Thy-1.1, CD45.2 and TCRvβ12 staining was analyzed by flow cytometry. Representative of three performed with similar results.

**Supplementary Figure 5:**
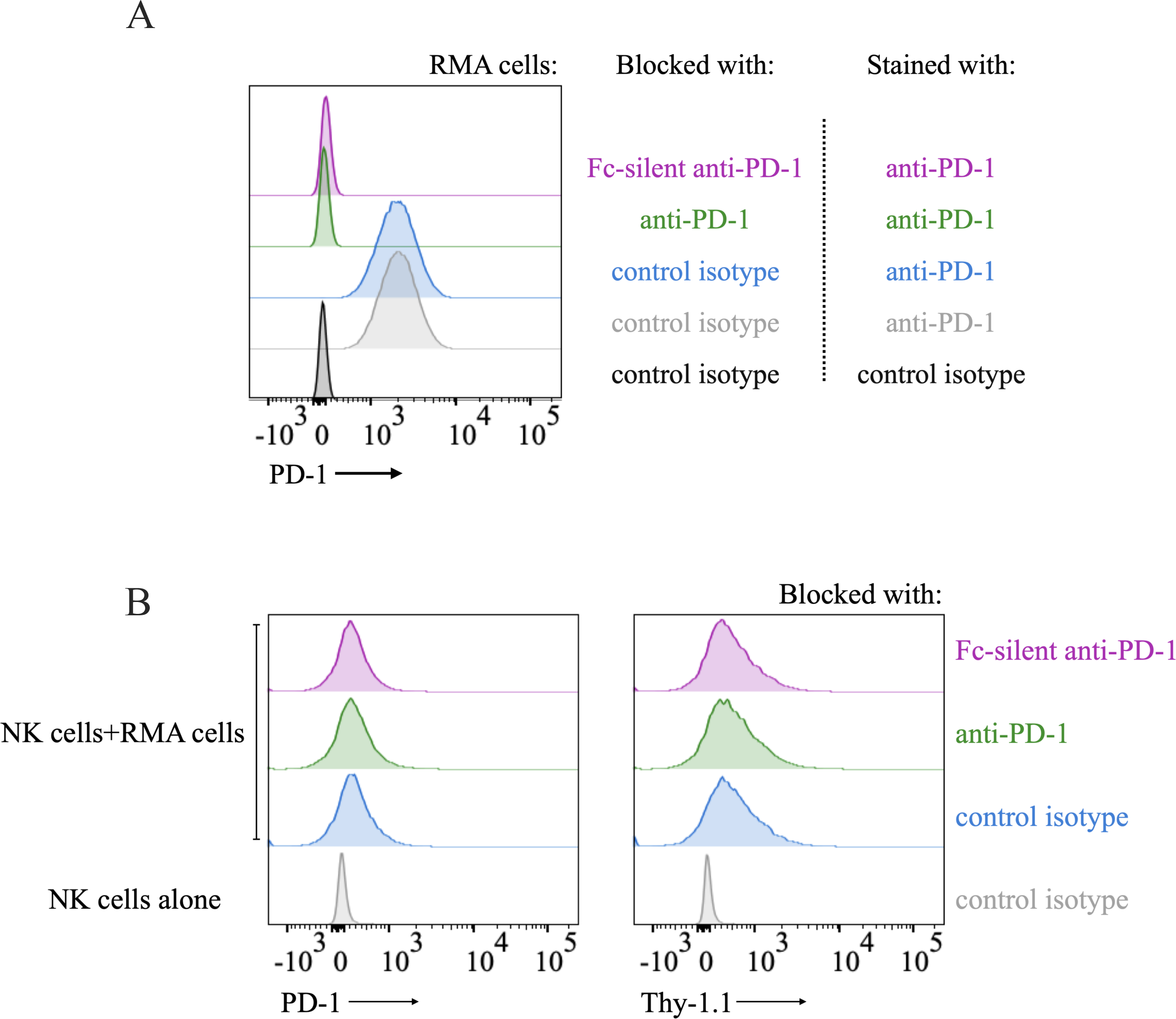
PD-1 antibodies do not affect or promote PD-1 trogocytosis by NK cells. (A) Saturation of PD-1 sites on RMA cells was assessed by stained with the PD-1 antibody cells that were previously co-incubated with anti-PD-1 or control isotype. (B) NK cells purified from *Pdcd1^-/-^* mice were incubated with RMA cells in the presence of the indicated blocking antibody and then stained for PD-1 and Thy-1.1. The experiment depicted is representative of three performed.

**Supplementary Figure 6:**
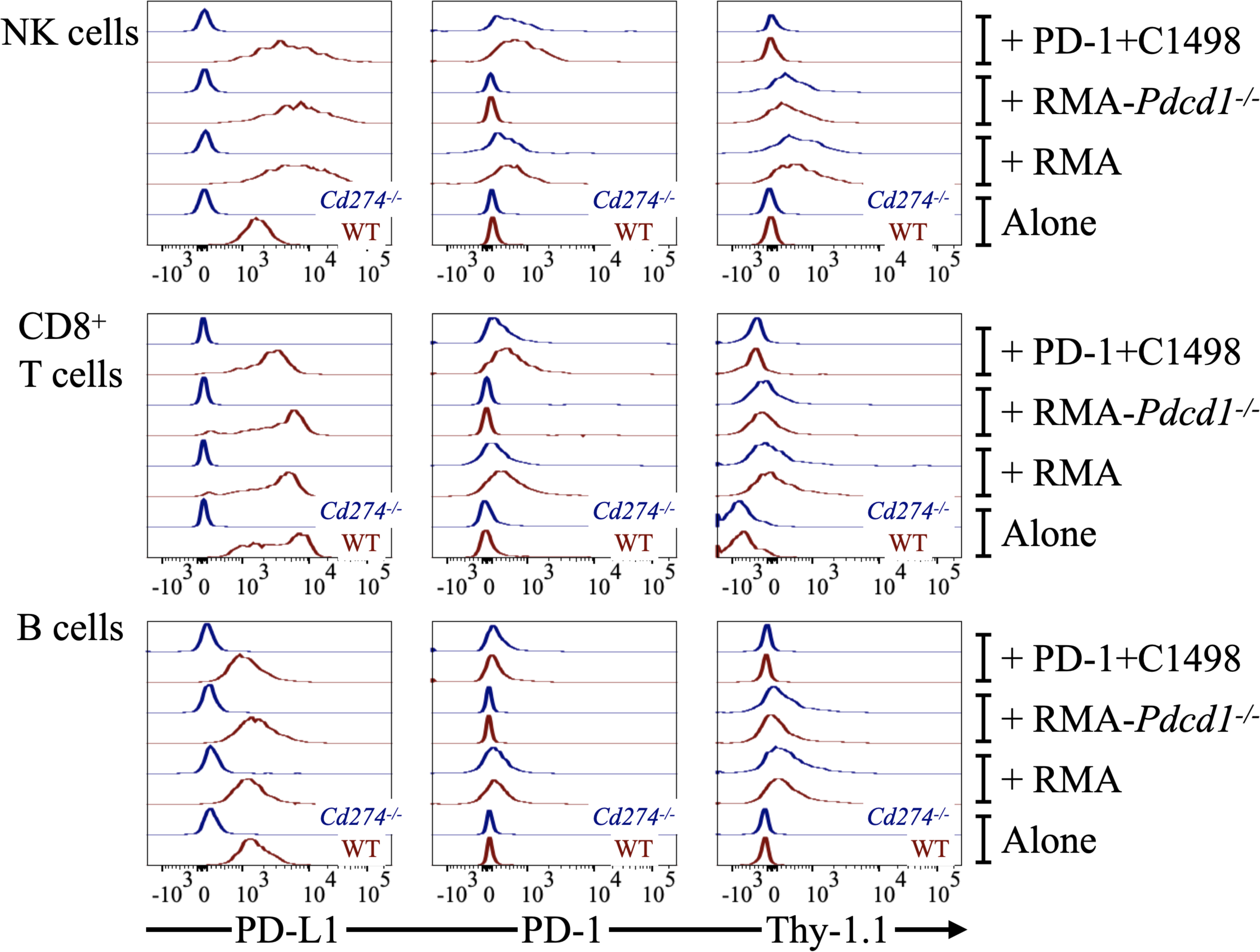
PD-L1 is dispensable for PD-1 trogocytosis in NK, T and B cells. *Cd274*^-/-^ or wild type splenocytes were cultured with tumor cells for 3 days before PD-1 and Thy-1.1 staining was assessed by flow cytometry. Representative of 3 mice/genotype analyzed.

**Supplementary Figure 7:**
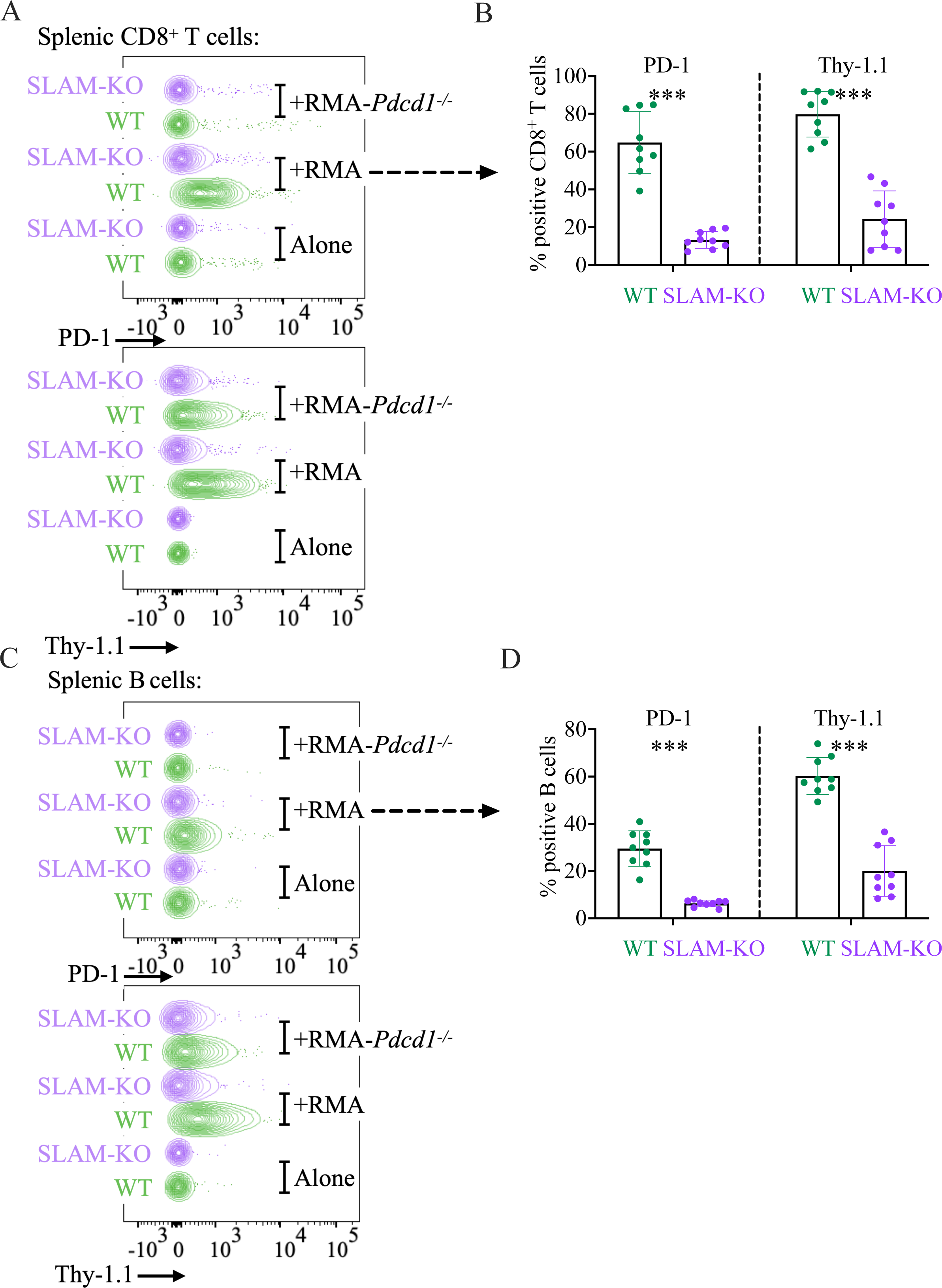
SLAM receptors are required for trogocytosis in CD8^+^ T and B cells. SLAM-ko or wild type splenocytes were cultured with tumor cells for three days. Representative staining and cumulative analysis are depicted. Statistical analysis with two-tailed unpaired Student’s t-test. ***: p<0.001.

**Supplementary Figure 8:**
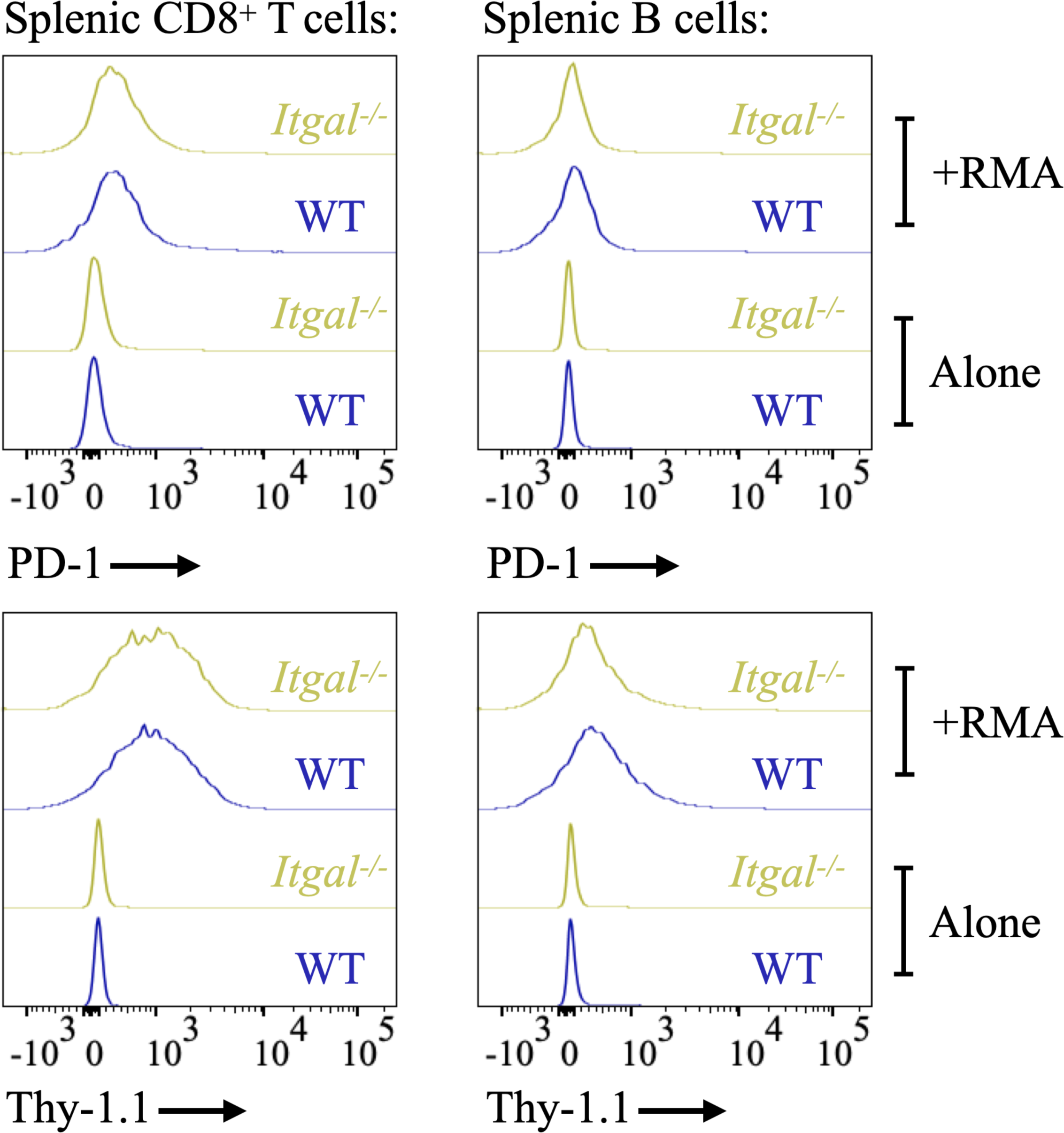
LFA-1 is not necessary for trogocytosis in CD8^+^ T and B cells. *Itga1^-/-^* or wild type splenocytes were cultured with tumor cells for three days when PD-1 and Thy-1.1 staining was assessed on CD8+ T cells and B cells.

**Supplementary Figure 9:**
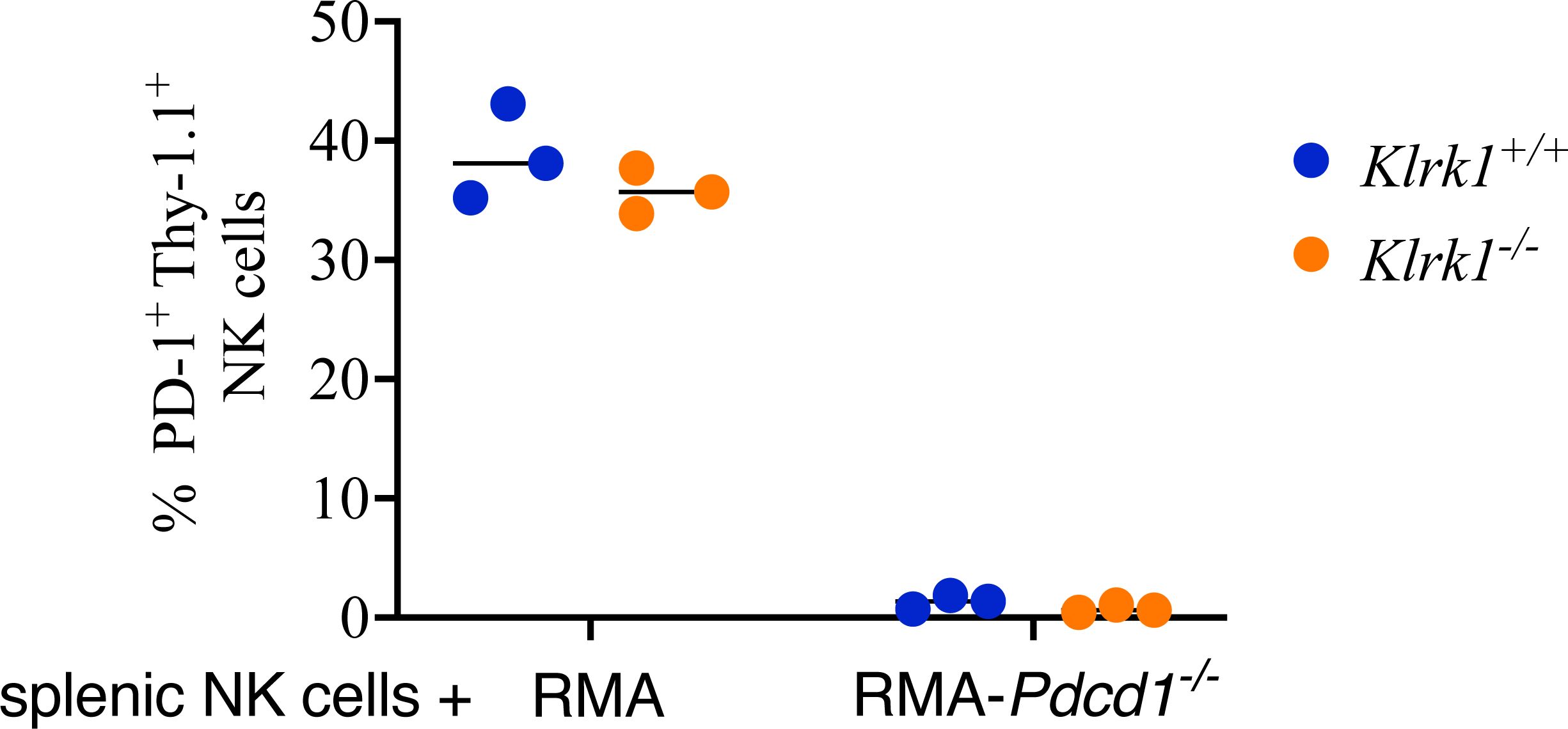
NKG2D is not required for trogocytosis in NK cells. *Klrk1^-/-^* or control littermates NK cells were cultured with tumor cells for three days and then stained for PD-1 and Thy-1.1.

**Supplementary Figure 10:**
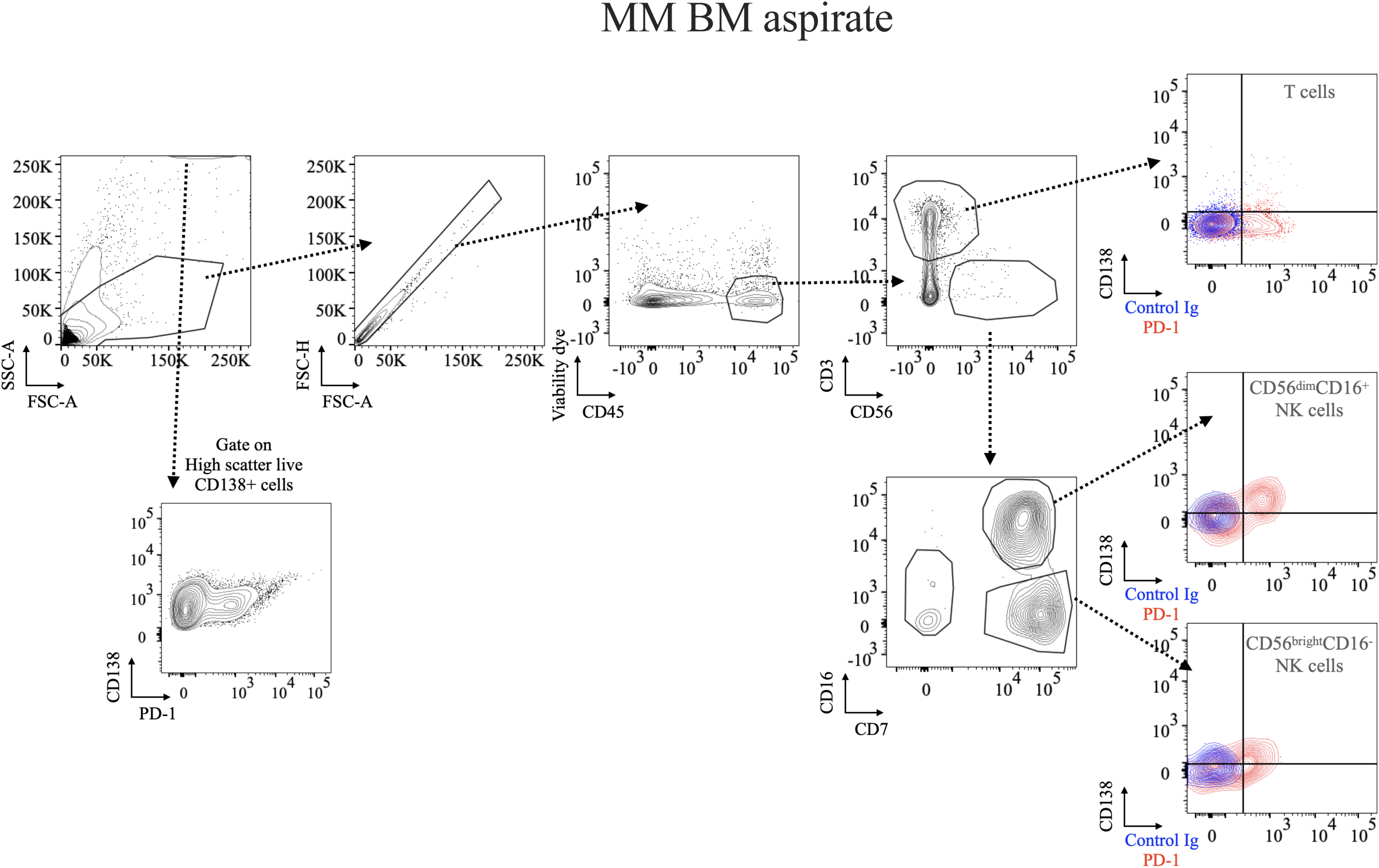
the gating strategy for NK and T cell identification in the bone marrow of patients with clonal plasma cell disorders is displayed.

